# Tumor Priming by Ultrasound Mechanogenetics for with SynNotch CAR T Therapy

**DOI:** 10.1101/2024.10.01.615989

**Authors:** Chi Woo Yoon, Chunyang Song, Dung Ngo Minh Nguyen, Linshan Zhu, Phuong Ho, Ziliang Huang, Gengxi Lu, Ali Zamat, Alexa Lewis, Ruimin Chen, Yushun Zeng, Nan Sook Lee, Christina Jamieson, K. Kirk Shung, Qifa Zhou, Yingxiao Wang

## Abstract

Cell-based cancer immunotherapy holds potential as a therapeutic approach, yet its application for solid tumor treatment remains challenging. We created a system where focused ultrasound (FUS) is able to remotely stimulate gene expressions in a specific tissue area through mechanical induction, gated by a chemical inducer to minimize the background noise. This system, known as CaDox, integrates FUS-triggered mechanical and calcium stimulation with doxycycline-responsive genetic circuits, which allows the localized expression of the clinically validated and specific antigen CD19 within a subpopulation of cancer cells upon FUS stimulation. These CD19-expressing cells can then function as “training centers” that activate synNotch chimeric antigen receptor (CAR) T cells to generate CARs that recognize a less specific but widespread antigen in cancer cells, thereby attacking and suppress the whole cancer cell population nearby at the tumor site. We validated the functionality of this CaDox system *in vitro*, in organoids, and *in vivo*, demonstrating its potential for various cell types and as a versatile platform for precisely controllable immunotherapy. Our combinatorial approach thus offers a FUS-controlled remote and non-invasive priming of solid tumors for effective and safe CAR T immunotherapy via the induced production of clinically validated antigens.

## Introduction

The ability to remotely control genetics and cellular functions with high spatiotemporal precision has long been a goal in the field of biomedicine. A variety of techniques have been developed to perturb cells and tissues through energy-based modalities, such as optics^1^, magnetics^2^, and ultrasound^3^. Among the listed modalities, optogenetics has emerged as a particularly successful approach, combining molecular sensors of lights and inducible gene expression systems to control protein production and cellular functions^4–9^. Optogenetics involves the use of light-sensitive proteins to manipulate cellular activity in response to specific wavelengths of light^1^. This powerful tool has enabled researchers to control various aspects of cellular function with unprecedented precision, both spatially and temporally. The versatility of optogenetics has led to its widespread adoption across various disciplines, including neuroscience^1, 9, 10^, immunology^11–13^, and developmental biology^14^, among others. However, a major challenge in translating optogenetics to clinical applications is the limited penetration depth of light, which restricts its utility in deeper tissues and organs.

Ultrasound is a promising external modulator for regulating cellular processes, as it is a form of mechanical energy that can be transmitted efficiently through deep tissues, offering several orders higher in penetration depth than that of light^15^. Previously, ultrasound has been recognized for its ability to activate genes indirectly via thermo effects^16–18^. However, the burgeoning understanding of mechanotransduction pathways in cells^19^, opens up a new perspective on ultrasound’s potential, suggesting a more prompt and direct approach for remote control of genetics and cellular behaviors with a higher precision in space^20–22^. We previously developed mechanogenetics, in which focused ultrasound (FUS) is used to activate specific mechanosensitive channels, Piezo 1, in cells physically coupled to microbubbles^23^. However, the *in vivo* application of this approach is limited by the uncontrollability of this microbubble co-factor, together with its large size^24^ and relatively short circulation time in the body^25^. Thus, there remains an unmet need for FUS-based mechanogenetics that can achieve genetic and cellular control directly via mechanical perturbation without the need of cofactors such as microbubbles.

The mechano-sensitivity of cancer cells has recently emerged as a critical character^26–28^. These cells can sense and respond to the physical properties of their surrounding extracellular matrix (ECM), such as stiffness, topography, and composition^29–31^. Interestingly, several studies have reported that some cancer cells are sensitive to FUS stimulation^32, 33^, suggesting that this modality could be harnessed to target and manipulate the genetics of these tumor cells in preparation for subsequent therapy.

Chimeric antigen receptor (CAR) T-cell therapy is a groundbreaking immunotherapy that involves the genetic modification of a patient’s T-cells to express CARs, which can specifically target and eliminate cancer cells^34^. This approach has shown remarkable success in treating hematological malignancies, revolutionizing cancer therapy. However, the application of CAR T-cell therapy in solid tumors faces major obstacles, including the challenge posed by the heterogeneity of solid tumors, which complicates the discovery of truly tumor-specific and homogenous antigens and makes it difficult to avoid life-threatening on-target off-tumor toxicity^35^. The remote and noninvasive control of genetics and cell activity with a high spatiotemporal precision would be an ideal solution to precisely address this issue, inducing the cancer killing actions at confined and desired tumor areas. As such, FUS-based mechanogenetics holds great promise as a potential candidate for achieving this level of control, providing a more spatiotemporally targeted and adaptable approach for solid tumor treatment.

In this study, we present a FUS-based mechanogenetics approach without any cofactor to remotely and noninvasively control designed cellular genetics, leveraging the mechanosensitivity of cancer cells in calcium signaling and a Dox-gated AND logic genetic circuit. We characterized this system to produce clinically validated antigen in a subpopulation of cancer cells, which served as “training centers” to engage and activate SynNotch CAR T cells to eradicate the whole cancer population in the tumor regions via a homologous antigen. As such, our technology provides a FUS-controlled tumor priming in genetics with clinically validated antigen for safe and effective CAR T immunotherapy.

## Results

### Mechanical induction by FUS of intracellular calcium responses in prostate cancer PC-3 cells

Our first goal was to identify a cancer cell that is sensitive to mechanical perturbation by short pulsed acoustic waves without microbubbles or any other co-factor, and exhibits robust intracellular calcium responses. We selected prostate cancer PC-3 cells for our study, as their sensitivity in calcium signaling to focused ultrasound (FUS) stimulation has been reported previously^33, 36^. To assess the FUS-mediated intracellular calcium response, we integrated an ultrasound stimulation system with an epi-fluorescence microscope (Fig. 1A). For the stimulation, we first employed a single element 35-MHz FUS transducer with parameters (see Methods) aiming to elicit a strong calcium response that could subsequently drive strong gene expressions. This transducer, operating within the low-intensity pulsed ultrasound (LIPU) range, delivers focused energy to a localized region measuring 120 µm in diameter (see supplementary Fig. 1A). The selected center frequency and output powers were chosen to avoid cell toxicity, consistent with previously reported conditions^33, 37^.

**Figure 1:**
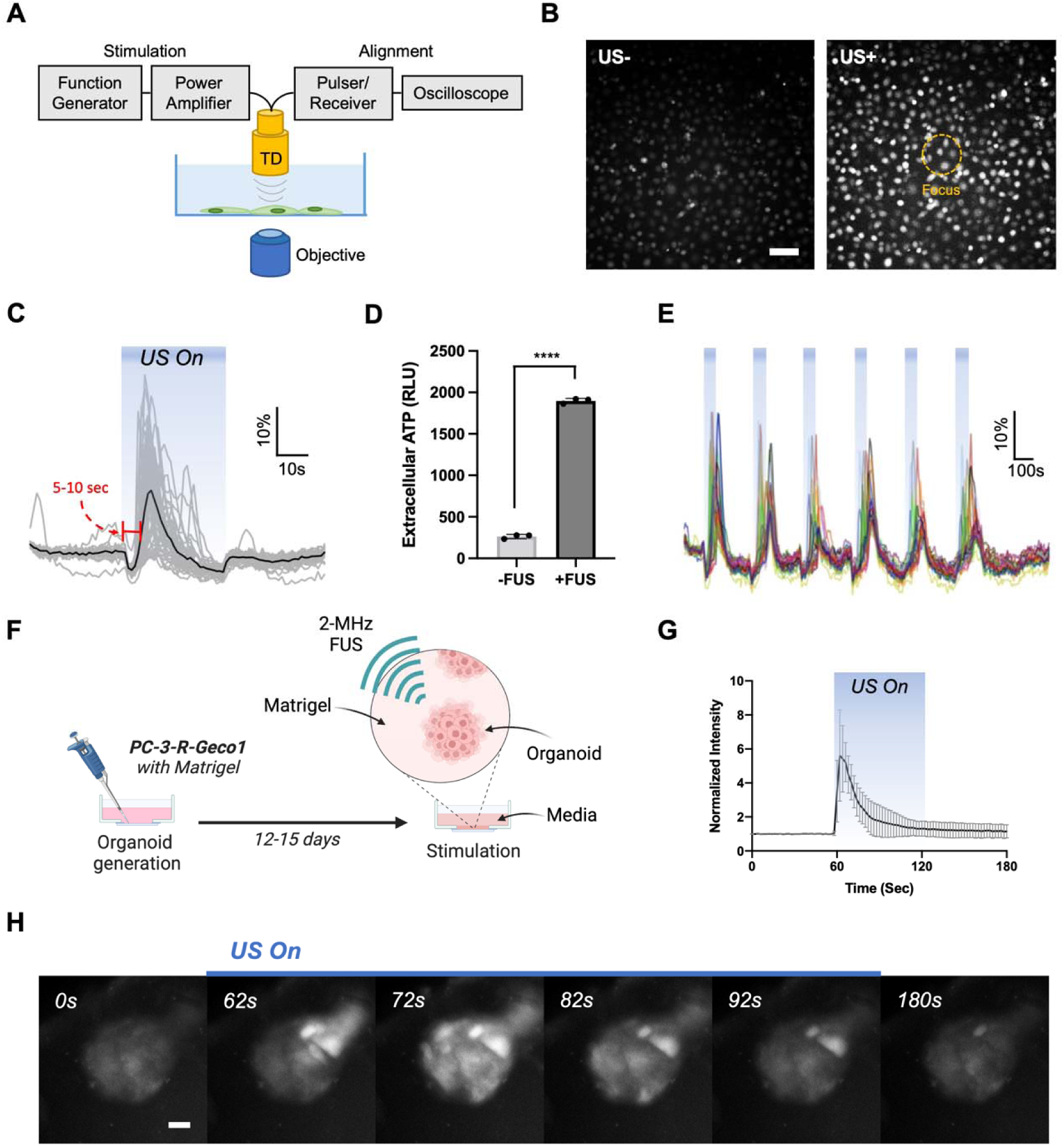
FUS induced calcium responses in prostate cancer PC-3 cells. (A) Schematic diagram of the FUS stimulation apparatus integrated into an inverted epifluorescence microscope to monitor real-time calcium dynamics upon FUS stimulation. (B) Representative images of intracellular calcium responses in PC-3 cells loaded with Fluo-4 am upon FUS stimulation. The orange dotted circle indicates the focal zone (120 µm) of the transducer. Scale bar: 100 µm. (C) The time courses of intracellular calcium levels within each cell body at the FUS focal zone (Grey lines), with the black line representing the average. (D) Luciferase-based ATP measurements from the imaging media with and without ultrasound exposure. n=3. (E) Calcium responses from short pulsed repeats of FUS stimulation: 1 min every 5 min for six repeats. (F) Schematic drawing of the organoid setup on a culture dish. Organoids were generated with PC-3 cells stably expressing R-GECO1. (G) Calcium response profiles of the organoid cells in response to 2-MHz FUS stimulation. FUS stimulation was on from 60 sec to 120 sec. Error bars represent SEM. (H) Representative time-lapse images of calcium responses from PC-3-derived organoids. Scale bar: 50 µm.

As anticipated, we observed strong calcium reactions in response to FUS stimulation (Fig. 1B, 1C). Interestingly, the cells located outside of the focal area of the FUS also showed responses through calcium propagation (Supplementary Fig. 2). We also observed a response latency of several seconds from the onset of the FUS stimulation (Fig. 1B). This response latency and the calcium propagation beyond the FUS-stimulated area can be attributed to the paracrine release of ATP by the mechanosensitive non-junctional hemichannel known as pannexin 1 (PANX1), as previously reported^36^. Upon activation, the mechanosensitive PANX1 can facilitate ATP release into the extracellular space^38^. The extracellular ATPs trigger the activation of purinergic receptors, leading to both calcium influx and intracellular calcium release in neighboring cells. As such, while FUS can directly stimulate ATP release through mechanical sensing, the overall calcium response is an indirect effect, mediated by the subsequent purinergic signaling^36^. This highlights the intricate interplay between mechanical and biochemical signaling pathways in FUS-induced cellular responses. Indeed, we detected a significantly increased level of extracellular ATP from the dish with FUS exposure (Fig. 1D). We further revealed that repetitive FUS stimulations (1 minute every 5 minutes for 30 minutes) could consistently elicit multiple calcium responses from PC-3 cells (Fig. 1E).

We further investigated these FUS responses in a 3D culture model to mimic the physiologically relevant conditions. We generated a 3D organoid model based on PC-3 cells that stably express the genetically encoded calcium sensor R-Geco1 (Fig. 1F), following the previously reported methods for organoid generation^39^. To stimulate the organoids ranging in size from 200 to 600 µm, we employed a clinically relevant low-frequency FUS transducer, which offers a larger coverage area (2-MHz, ∼700 µm). The output power of this transducer was tuned to fall within the LIPU range to maintain the consistency of different experimental conditions (Supplementary Fig. 1B). We found that the PC-3 cells in organoids also responded to FUS with strong calcium signals (Fig. 1G-H). These calcium waves were similarly observed to spread from the stimulated area to neighboring cells, reinforcing the possible involvement of paracrine signaling of ATP. These findings suggest that the FUS-induced mechanical perturbation is sufficient to induce intracellular calcium responses in both 2D and 3D culture models.

### Development and characterization of AND logic gate genetic circuit: CaDox

We then explored the possibility of designing genetic transducers and circuits that can convert the accumulated effect of FUS-induced calcium dynamics into controlled gene expressions. Nuclear factor of activated T cells (NFAT) is a family of transcription factors that play important roles in regulating gene expression in immune cells as well as in other cell types, including cancer cells^40^. NFAT proteins can be activated by intracellular calcium signaling, and subsequently translocate to the nucleus to regulate the transcription of target genes. To investigate whether the FUS-mediated calcium responses in PC-3 cells can be sufficient to induce NFAT translocation, we introduced a lentiviral vector to express NFAT protein fused with EGFP in PC-3 cells. A clear translocation of NFAT proteins was observed upon repetitive FUS stimulations (Fig. 2A), suggesting that the NFAT translocation may be utilized to control transcriptional activity upon FUS-mediated calcium.

**Figure 2:**
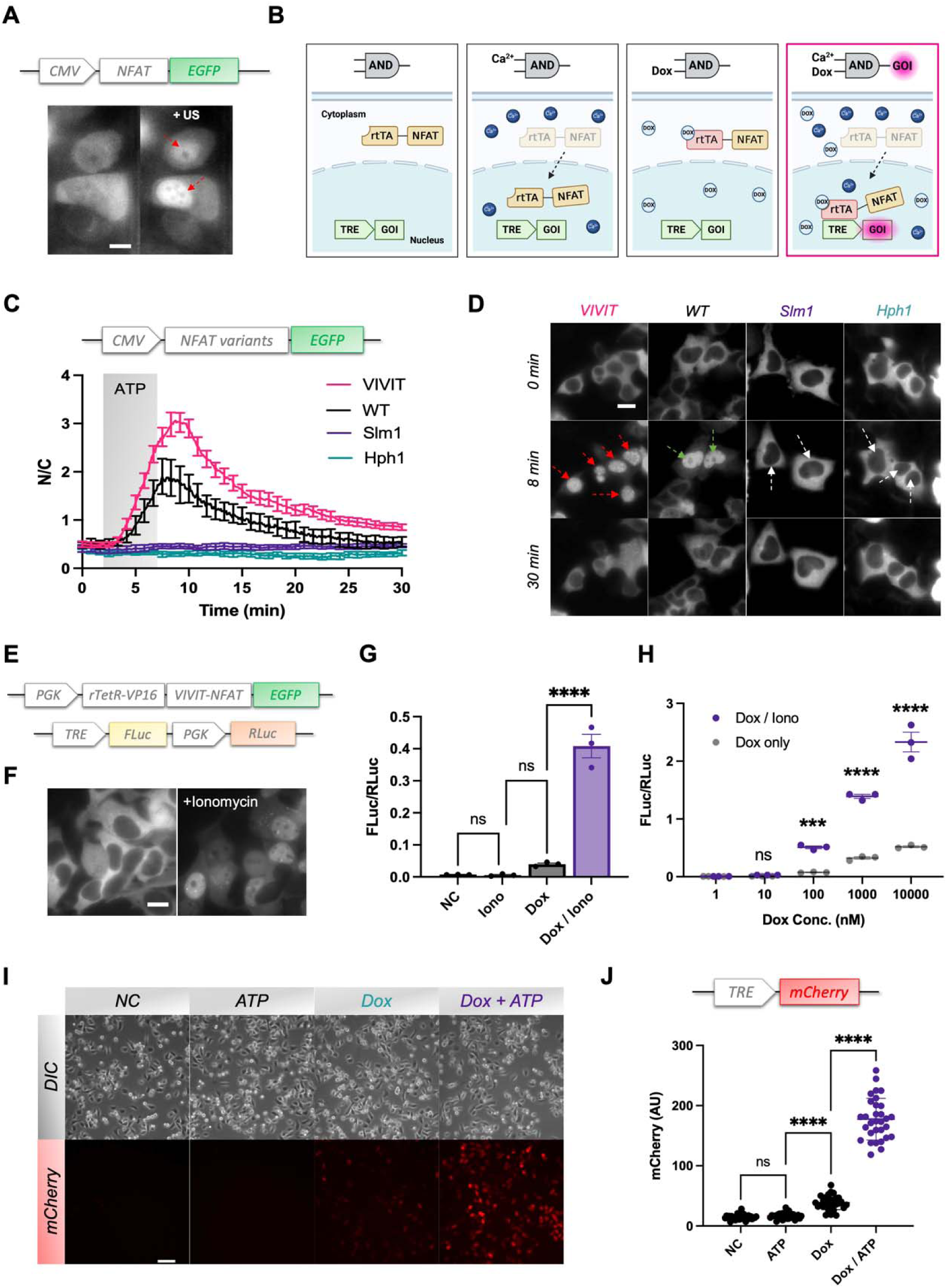
Development and characterization of the CaDox system. (A) NFAT translocation upon repetitive FUS stimulations tracked by EGFP directly fused to NFAT. Scale bar: 10 µm. (B) Schematic diagram of the calcium- and doxycycline-gated genetic circuit, termed CaDox. (C-D) Translocation time courses (C) and images (D) of the NFAT variants with different affinities for calcineurin (CN). ATP (60 µM) was added at 2 min to induce calcium elevation and washed off at 7 min. n=5. (E) CaDox system consists of CaDox regulator and CaDox reporter. (F) The translocation of CaDox regulator upon ionomycin (1µM) induction. (G) Gene duction quantification of the CaDox system using firefly luciferase (Fluc) as an inducible reporter and renilla luciferase (RLuc) as a constitutive reference control. n=3. (H) Titration of doxycycline concentration with and without ionomycin. n=3. Statistical significance was determined by t-test. (I-J) Representative images (I) and quantifications (J) of CaDox system in PC-3 cells with mCherry as a reporter, under various treatments as indicated. Scale bar: 100 µm. n=31. Error bars represent SEM. Statistical significance was determined by ANOVA with Tukey’s multiple comparison test unless indicated otherwise.

Next, we aimed to develop a genetic transducing module that can transduce FUS-induced calcium signaling into user-defined gene expressions. We previously developed a genetic circuit coupling a minimal promoter with the NFAT response element (NFAT-RE), and two other calcium-dependent elements: the cAMP response element (CRE) and the serum response element (SRE), to transduce calcium signals into gene expressions^23^. However, besides the varying expression levels of endogenous NFAT in different individual cells^41^, this system can be vulnerable to spontaneous calcium noises in cancer cells^42^. We indeed found that a genetic transducer based on NFAT-RE had a strong background noise of genetic activities in PC-3 cells, reaching levels comparable to the calcium ionophore ionomycin-induced gene production monitored by a GFP reporter (Supplementary Fig. 3).

To mitigate the noisy calcium-mediated genetic background as well as the varying levels of endogenous NFAT activity in individual PC-3 cells, we decided to engineer a synthetic transcription activator (STA) based on reverse tetracycline transactivator (rtTA) and NFAT protein. rtTA can regulate gene expression by binding to the tetO DNA sequence, in the presence of the antibiotic tetracycline or its derivative doxycycline^43^. Thus, by directly fusing the rtTA with the NFAT, this STA functions as an AND logic circuit, requiring the presence of both calcium signaling for rtTA-NFAT translocation and doxycycline for STA/DNA interaction to initiate transcription (Fig. 2B).

We first improved the STA by engineering NFAT with enhanced shuttling efficiency, by modifying its calcineurin docking sites. Calcineurin is a calcium-dependent phosphatase that becomes activated when calcium levels increase, leading to NFAT dephosphorylation and translocation to the nucleus^44^. Thus, we hypothesized that by modifying the calcineurin binding sites in NFAT, we may alter its translocation efficiency. We selected several mutants of calcineurin docking sequence with different dissociation constants (Kd) (Supplementary Fig. 4A) and used them to engineer GFP-tagged NFAT variants to track their translocation upon ATP-induced calcium dynamics (Supplementary Fig. 4B). The imaging results showed that the binding affinities between NFAT and calcineurin can significantly affect the translocation of NFAT. The VIVIT variant, which had the lowest Kd (0.5 µM), exhibited the fastest and strongest translocation upon calcium increase induced by ATP (Fig. 2C-D). Interestingly, by slightly lowering the binding affinities (Kd: 25 uM to 40 uM), NFAT completely lost its shuttling capability upon calcium stimulation. This result showed that the engineered NFAT variants with better binding affinities toward calcineurin can enhance the translocation capability, which could potentially improve the efficacy of genetic and transcriptional regulation in response to FUS stimulation and calcium signaling.

We further truncated the DNA-binding domain (RHD) of NFAT to avoid the interference of endogenous NFAT pathways, while retaining the calcium-dependent translocation of NFAT. The truncated NFAT (amino acids 4-399) can indeed shuttle in and out of the nucleus with a high efficiency upon calcium signaling (Supplementary Fig. 4C). We then created the STA by fusing the truncated VIVIT-tNFAT (amino acids 4-399) with rtTA to enable the gating by doxycycline. The resulting STA was hence designed to have the gene expression outcome precisely controllable by the doxycycline dosage. We named this Calcium- and Doxycycline-gated STA as CaDox regulator, and its corresponding Tetracycline Response Element (TRE)-based reporter circuit as the CaDox reporter. Together, these components comprise the CaDox system (Fig. 2E).

We verified the calcium-dependent translocation of CaDox regulator by tracking EGFP (Fig. 2F), which showed efficient translocation upon calcium elevation induced by 1 µM ionomycin. We then examined whether CaDox system could be tuned to minimize the transcriptional leakage engendered from spontaneous calcium noise. To conduct gene expression assays, we established a dual luciferase reporter system with inducible firefly luciferase (FLuc) and constitutive renilla luciferase (RLuc) as an internal reference in HEK293T cells. The cells were treated with doxycycline and/or ionomycin for one hour, then washed before subjected to a luciferase assay six hours after the drug treatment. The results showed that the CaDox system had minimal leakage in the absence of doxycycline (Fig. 2G). While there was minor background noise when the cells were incubated with doxycycline only, significant gene inductions were only observed in the presence of both calcium and doxycycline (Fig. 2G). The level of induction could be precisely controlled by the dosage of doxycycline (Fig. 2H), allowing tunability to meet varying needs of different applications. Using mCherry as the reporter, the CaDox system also showed a gated induction by doxycycline and ATP-induced calcium in PC-3 cells (Fig. 2I-J). This CaDox system is applicable across different cell types, including Jurkat T cell line and MDA-MB-231 breast cancer cells (Supplementary Fig. 5). As such, we engineered an AND logic gene circuit, the CaDox system, which is applicable to induce an efficient calcium-mediated gene expression gated by doxycycline with minimal background noise.

### Integration of CaDox system with FUS for mechanogenetics (FUS-CaDox)

Next, we evaluated the functionality of the CaDox system in controlling genetics upon FUS stimulation. We engineered CaDox-PC-3 cell lines with mCherry as a reporter gene to generate 3D organoids^39, 45^, which were subjected to treatment with both doxycycline and FUS (Fig. 3A). Specifically, organoids were pre-incubated with 200 nM of doxycycline, followed by repetitive FUS stimulations for 30 min (2 min On and 3 min Off for 6 times) using a 2-MHz transducer (peak-negative-pressure: 0.63 MPa) with pulsed waves (duty cycle: 10%) to activate the CaDox system (Fig. 3A). The imaging results revealed that neither FUS nor doxycycline alone resulted in a significant gene induction (Fig. 3B, C). However, when both stimuli were applied, the CaDox system led to a clear and significant gene induction compared to all other groups (p<0.0001). These results showed that the mechanical stimulation by FUS can be transduced by the CaDox system to control designed gene expressions in cancer cells.

**Figure 3:**
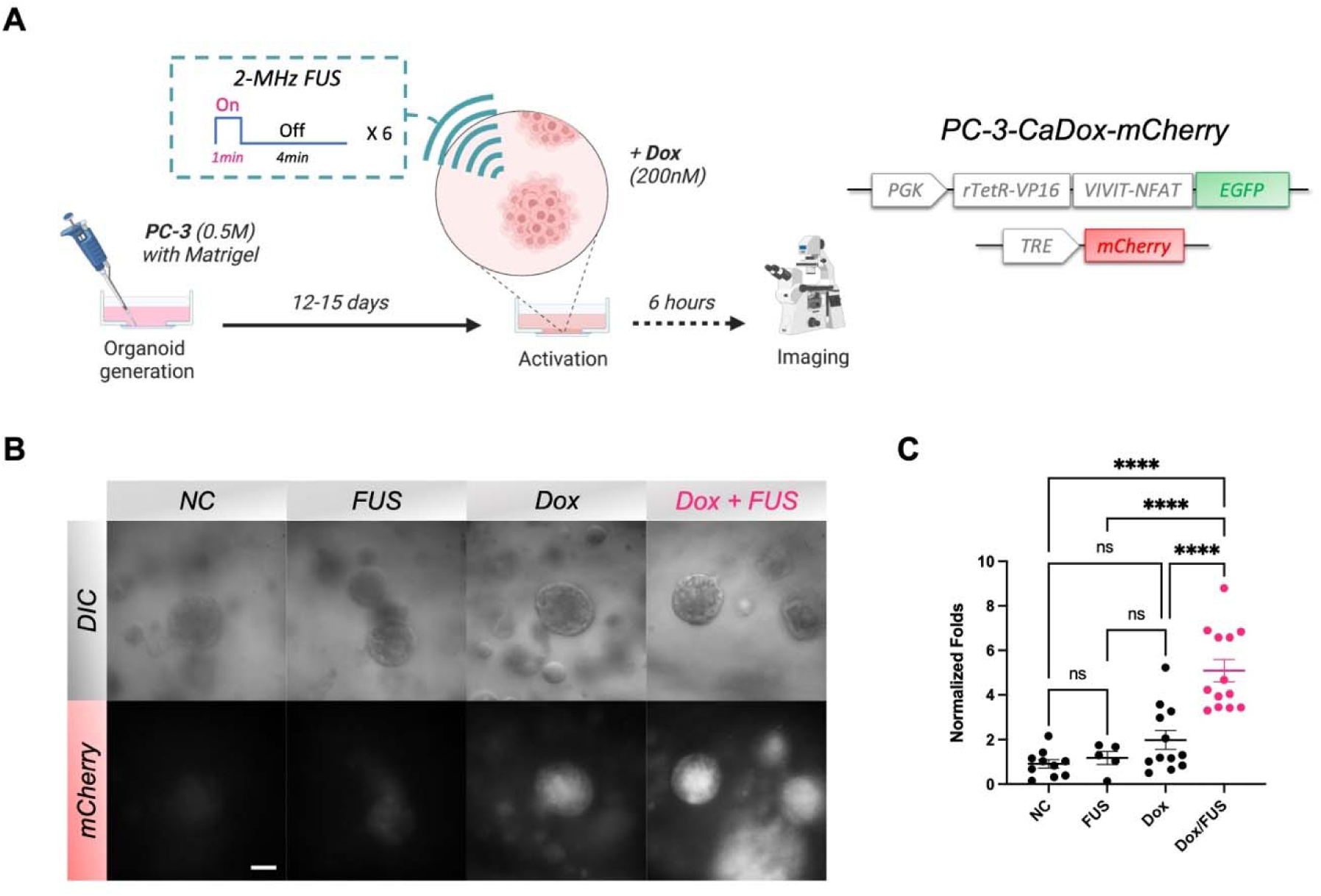
The FUS-based mechanogenetic system in the 3D organoid model. (A) Schematic diagram of organoid generation, activation, and imaging. PC-3-CaDox-mCherry cells were used to generate organoids. (B-C) Representative images (B) and quantifications (C) of organoids derived from PC-3 cells stably expressing the CaDox-mCherry system, under various treatments as indicated: the organoid bodies upon no treatment (n=10), FUS-only (n=5), doxycycline-only (n=12), and combined treatment groups (n=13). Scale bar = 50 µm. Error bars represent SEM. Statistical significance was determined by ANOVA with Tukey’s multiple comparison test.

### *In vivo* characterization of FUS-CaDox and mechanogenetics

To examine the FUS-CaDox system and mechanogenetics *in vivo*, we performed FUS stimulation experiments using engineered PC-3-CaDox cells in a xenograft mouse model. We designed a custom FUS stimulation apparatus in-house (Fig. 4A, Supplementary Fig. 6) featuring a 1-MHz focused transducer (diameter: 70 mm, radius-of-curvature: 65 mm) to deliver targeted mechanical stimulation to a localized volume at the tumor site (Fig. 4B, Supplementary Fig. 7). The FUS parameters are chosen to generate significant mechanical perturbations while maintaining safety thresholds and avoiding non-specific tissue disruptions. As such, the FUS power (peak-negative-pressure: 1.23 MPa) and repetition frequency (1-minute intervals every 2 minutes over a 30-minute period, Fig. 4A) were employed to maintain the mechanical index below the FDA safety guideline of 1.9 and avoid the risk of cavitation effects^46^. This setting also allows us to control the thermal output of the system so that the temperature variation did not exceed 1 (Fig. 4C), circumventing potential non-selective thermal effects that could confound our results. Together, this approach allowed us to precisely explore the genetic control of mechanical stimulation using FUS in an *in vivo* setting while adhering to safety guidelines.

**Figure 4:**
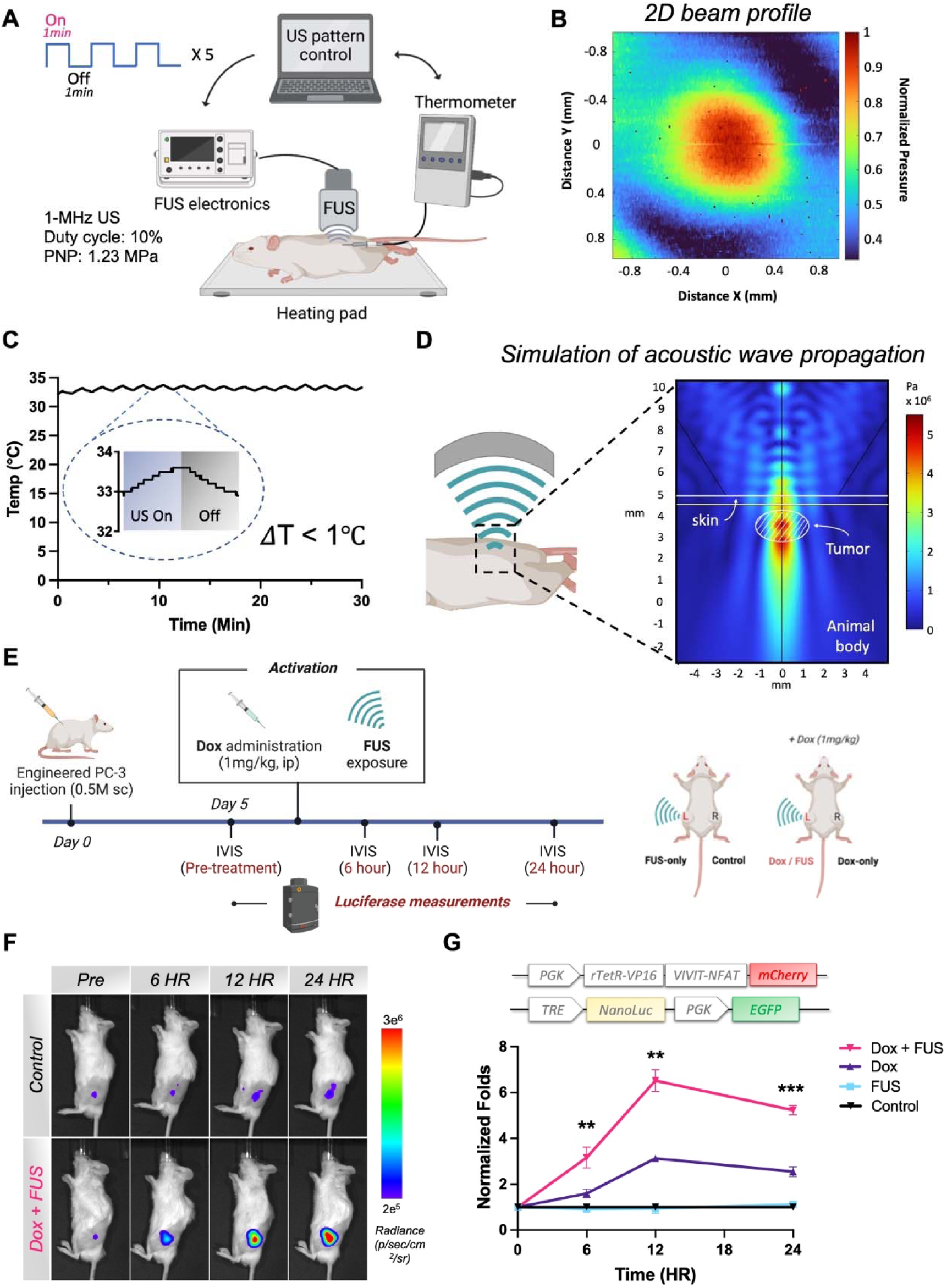
*In vivo* characterization of the FUS-based mechanogenetic system. (A) Schematic diagram of the custom-built FUS stimulation system for small animals. User-defined stimulation patterns and ultrasound parameters were input using a Python-based code that controls FUS electronics, a function generator, and a power amplifier, to send signals to the FUS transducer. Temperature changes in the focused area were measured by a needle-type thermometer during the 30-min stimulation window. (B) Two-dimension beam profile of 1-MHz transducer at the focus as measured by a hydrophone. (C) Temperature readings from the thermometer located at tumor indicating less than 1℃ of temperature fluctuation during FUS stimulation. (D) Simulation results of acoustic wave propagation along the wave path and mechanical stress distribution in the animal body. In the simulation, an animal body environment was created with a skin layer (0.5 mm thickness), a muscle layer, and a 3 mm x 3 mm x 1.8 mm (L x W x H) tumor embedded within the muscle layer. (E) Experimental design of *in vivo* activation of the CaDox system by FUS. PC-3-CaDox-NLuc cells were injected into NSG mice to establish a bilateral tumor model. On Day 5, tumors were treated with either doxycycline and/or FUS. Bioluminescence imaging (BLI) was utilized to monitor luciferase intensities over a 24-hour period. (F) Representative BLI images of gene expression in subcutaneous tumors kept untreated, or in response to the treatment with the combined doxycycline and FUS. (G) Normalized fold changes in gene expression of subcutaneous tumors over 24-hours under different treatments (n=4). The gene expression of each treated condition was normalized against that of the untreated group for accurate comparisons. Error bars represent SEM. Statistical significance was determined by paired t-test. Comparisons between treatments of doxycycline-only and combined doxycycline and FUS yielded p-values of 0.0098 (6-HR), 0.0033 (12-HR), and 0.0003 (24-HR).

To gain a more comprehensive understanding of acoustic wave propagation within the animal body and to estimate the amount of energy deposited at the focus of the ultrasound, we conducted simulations using COMSOL Multiphysics to assess the spatial distribution of acoustic wave pressure^47, 48^. Our findings revealed a minimal difference in the pressure peak and distribution of the focused ultrasound, when accounting for the attenuation due to the animal’s skin and muscle (Supplementary Fig. 8)^47, 48^. Our results also demonstrated that FUS can generate a focal pressure distribution covering an area of approximately 1.5 mm in the X-Y direction and 2 mm in the Z direction (Fig. 4D).

We next engineered CaDox-PC-3 cell lines with an inducible Nano luciferase (NLuc) reporter and implanted them into NOD scid gamma (NSG) mice to establish a bilateral tumor model (Fig. 4E). Five days post-implantation, we divided the mice into two groups. One group received doxycycline (1 mg/kg) while the other group was left untreated. FUS stimulation was then applied to the left tumors of mice in both groups, leaving the right tumors as unstimulated controls for reference. We monitored the NLuc expression levels before treatment, as well as 6-, 12-, and 24-hours post-doxycycline administration, to track the temporal gene induction dynamics of the CaDox system activation.

Significant gene expression was observed only in the group receiving the combination of doxycycline and FUS treatment (left tumor) (Fig. 4F, G, and Supplementary Fig. 9). As expected, no induction of gene expression was observed in the non-treated control group, nor the group with FUS stimulation alone, in the absence of doxycycline. A minor and statistically insignificant induction of gene expression can be observed in the control group with Dox only treatment. No signs of tissue damage or alteration in tumor growth were observed at the FUS-treated sites. These results underscore the safety and applicability of the FUS-CaDox system for *in vivo* control of designed gene expressions.

### Mechanogenetics-based combinatorial immunotherapy

We then applied our FUS-CaDox mechanogenetic system for tumor priming in immunotherapy, integrating with synNotch and chimeric antigen receptor (CAR) T cells^49^. FUS-CaDox is rewired to produce a clinically validated and specific antigen CD19 upon the Dox-gated FUS stimulation. These induced subpopulation of tumor cells expressing CD19 can then serve as priming cells and “training centers” to stimulate the subsequently infused CD19-synNotch CAR T cells, which can be activated through a synthetic Notch receptor recognizing CD19 to produce a CAR against a less specific but widespread tumor antigen expressed on tumor cells to eradicate the whole tumor population at the tumor site and its neighboring regions (Fig. 5A). Therefore, the FUS-CaDox priming of tumor cells integrated with the killing by synNotch CAR T cells can overcome the problems: (1) the lack of clinically validated antigens for solid tumors; (2) the low gene delivery efficiency *in vivo*; and (3) the limitations in spatial coverage of FUS.

**Figure 5:**
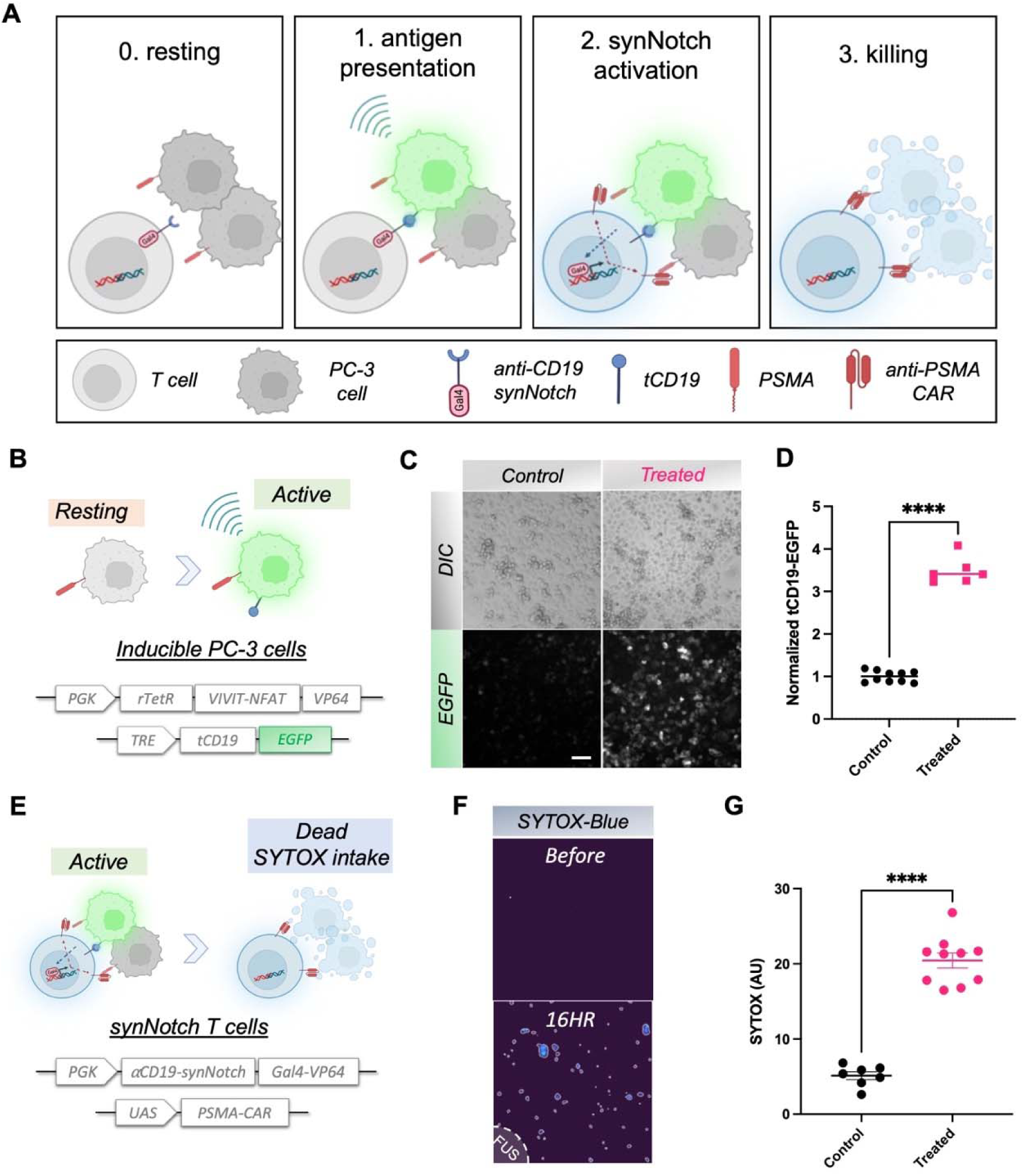
Combinatorial immunotherapy *in vitro*. (A) Schematic diagram of the combinatorial immunotherapy approach. The subsets of tumor cells expressing tCD19 upon FUS activation can guide synNotch CAR T cells to target the whole population of cancer cells at the tumor site, via a CAR produced to recognize a homologous antigen PSMA. (B) PC-3 cells were equipped with the CaDox-tCD19-EGFP system to express tCD19 and EGFP upon FUS activation. (C-D) Representative images of induced gene expression (C) and Quantified EGFP expression levels measured (D) in no treatment (n=10), or combined treatment groups (n=6). (E) Primary T cells were engineered with anti-CD19 synNotch and inducible anti-PSMA-CAR to be activated by the induced tCD19^+^ cells to attack and clear PSMA^+^ cancer cells. (F) Representative images of SYTOX staining to assess the cytotoxicity of PC-3 cells. (G) Quantification of cytotoxicity of PC-3 cells in no treatment (n=7), and combined treatment groups (n=10). Error bars represent SEM. Statistical significance was determined by ANOVA with Tukey’s multiple comparison test.

To validate the concept of the combinatorial immunotherapy, we first generated PC-3-CaDox cell lines with the CaDox reporter encoding an inducible truncated CD19 (tCD19), which has been clinically validated to serve as a specific and efficient target for CAR T therapy (Fig. 5B)^50^. We fused EGFP to the cytoplasmic-tail of tCD19 and verified that tCD19 retains plasma membrane localization by imaging, and confirmed that the extracellular epitope of tCD19 can be recognized by anti-CD19 antibody (Supplementary Fig. 10). Consistent with our previous results (Figs. 4F-G), a combined treatment of doxycycline and FUS triggered significant tCD19 expression on the cell membrane (Fig. 5C, D, and Supplementary Fig. 11).

After confirming the tCD19 inducibility of the cancer cells, we genetically engineered Jurkat T cells with an anti-CD19 synNotch receptor to drive tagBFP as an activation reporter for synNotch CAR T (Supplementary Fig. 12A). The activation of Jurkat T cells, as indicated by tagBFP, was significantly higher when they were co-cultured for 24 hours with PC-3 cells pre-activated by the Dox-gated FUS to express tCD19 (Supplementary Fig. 12B). This result indicates that tCD19 induced by FUS-CaDox can train and lead to synNotch activation and reporter production in T cells.

We then examined the synNotch-mediated killing using primary human T cells. For the homologous antigen broadly expressed in prostate cancer cells, we chose prostate-specific membrane antigen (PSMA). We hence introduced the inducible anti-PSMA CAR in the synNotch CAR T cells (Fig. 5E). To characterize the cytotoxicity and specificity of the CD19/synNotch-PSMA/CAR (synNotch-CAR) T cells, we first generated three different PC-3 cell lines, each constitutively expressing either tCD19, PSMA, or both tCD19 and PSMA. Each of these three PC-3 cell lines was co-cultured with the synNotch-CAR T cells (E:T = 1:1) for 24 hours before assessing the killing efficiency. As expected, without the PSMA antigen, no significant cytotoxicity was observed (Supplementary Fig. 13A). A minor cytotoxicity was observed in the CD19^-^/PSMA^+^ group (p-value: 0.0419), but significant killing was only found in the CD19^+^/PSMA^+^ group (p-value < 0.0001, Supplementary Fig. 13A), demonstrating that our synNotch-CAR T cells can specifically recognize and kill cells expressing both CD19 and PSMA. Next, we examined whether the expression of tCD19 on a subpopulation of cancer cells could lead to the killing of the whole tumor population with a homologous PSMA antigen. We activated the CaDox-inducible tCD19 in PSMA-expressing PC-3 cells to achieve an approximate 15% tCD19^+^ subpopulation (Supplementary Fig. 13B). When we co-cultured these mixed tCD19^+^/^-^ cells with synNotch-CAR T cells, the fraction of viable cells dropped to 0.56 within 24 hours and 0.24 within 48 hours at an E:T ratio of 1:1, in contrast to the non-affected vial fraction of pure CD19^-^ PSMA^+^ PC-3 cells (Supplementary Fig. 13B). This result suggests that once trained and activated by the inducible tCD19, synNotch-CAR T cells can kill adjacent tumor cells expressing PSMA, no longer dependent on tCD19 anymore^49, 51^. SYTOX-blue staining consistently showed a significant killing of the cancer cells in the neighboring regions in and outside of the area exposed to Dox-gated FUS stimulation (Fig. 5F-G).

### *in vivo* examination of combinatorial cell-based immunotherapy

Next, we examined the tumor priming effect of FUS-CaDox mechanogenetic system integrated with synNotch-CAR T cells for cancer immunotherapy in an *in vivo* model. We followed the same experimental design and FUS parameters that we used for the *in vivo* FUS-CaDox gene inducibility testing (Fig. 4). Specifically, we first injected NSG mice on each side of the mice with 0.5 million PSMA^+^ PC-3 cells expressing inducible CaDox-tCD19 to establish bilateral tumor models. We continuously monitored tumor growth using bioluminescence measurements starting from Day 5. On Day 10, we separated the mice into two groups, with or without the intraperitoneal doxycycline administration (1 mg/kg) and stimulated all mice on the left tumors only with FUS (Fig. 6A).

**Figure 6:**
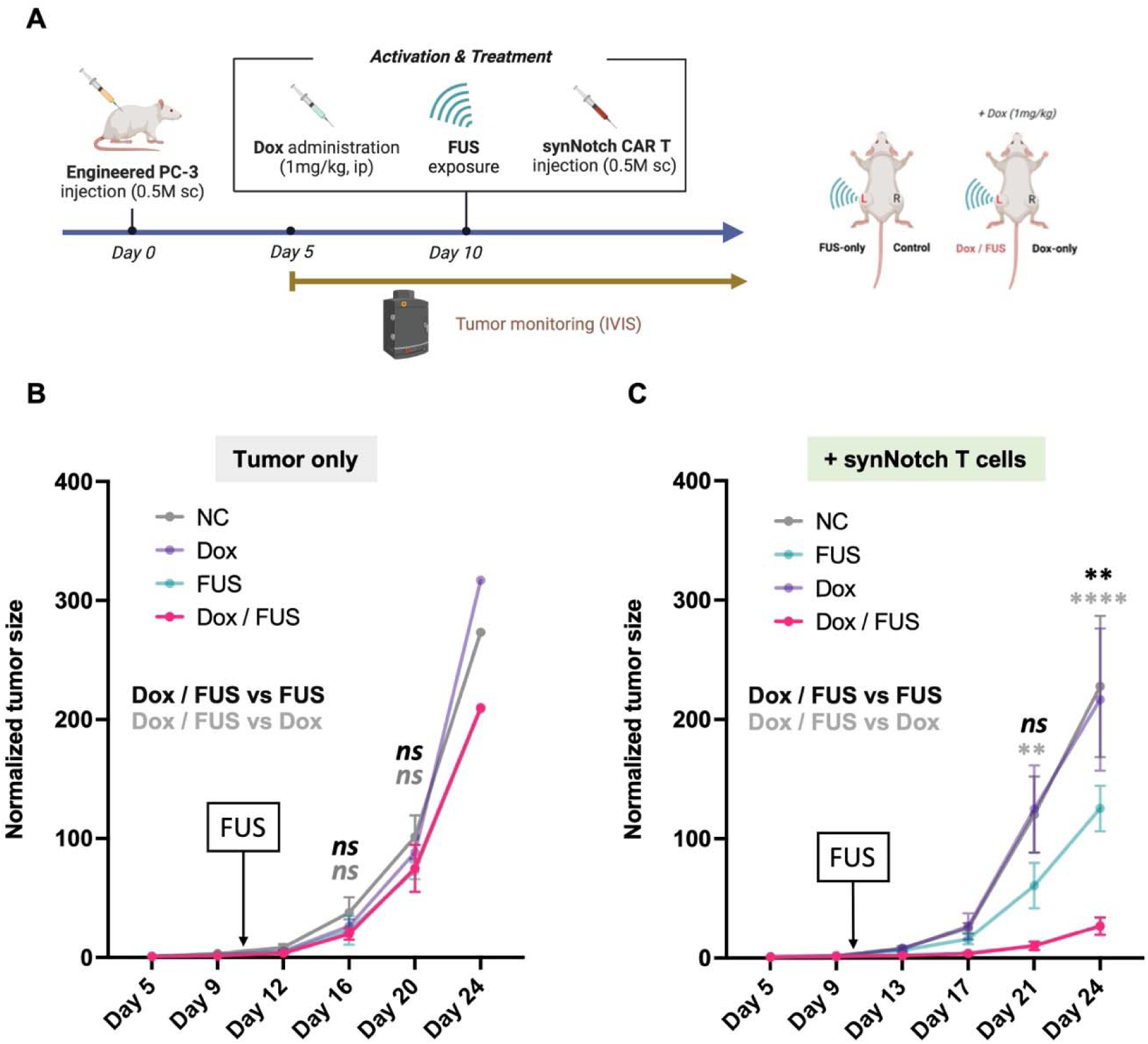
*In vivo* immunotherapy. (A) Schematics of the experimental timeline. Bilateral tumors were established by subcutaneously injecting engineered PC-3 cells on both sides of the NSG mice. (B) Quantification of normalized tumor growth after FUS-CaDox priming but without synNotch T cells. (n=4) (C) Quantification of normalized tumor growth after FUS-CaDox priming followed by synNotch T cells across different treatment groups as indicated. (n=5) Error bars represent SEM. Statistical significance was determined by ANOVA with Tukey’s multiple comparison test.

We showed there was no statistical difference between the FUS-CaDox primed and control groups on tumor growth (Fig. 6B, Supplementary Fig. 14). We then administered subcutaneously into each tumor site 0.5 million synNotch-CAR T cells right after the FUS treatment to assess the priming effect on tumor growth (Day 10, Fig. 6A). Tumor growth in the doxycycline- or FUS-only group was not statistically different from that of the negative control group without any treatment (Fig. 6C, Supplementary Fig. 15). In contrast, synNotch-CAR T cells can significantly suppress the FUS-CaDox primed tumor growth (Fig. 6C), demonstrating the effectiveness of our FUS mechanogenetic priming, in combination with synNotch-CAR, to treat solid tumors *in vivo*.

## Discussion

In this study, we developed a FUS-based mechanogenetics approach without the need of co-factors such as microbubbles, aiming to reveal the translational potential of direct, remote and non-invasive control of cellular mechanics and designed genetics using ultrasound. To achieve this, we exploited the mechanosensitivity of cancer cells and rewired the FUS-mediated calcium responses to user-defined transcriptional activity using the calcium and doxycycline-dependent AND logic gated circuit, referred to as the CaDox system. This approach allowed us to apply FUS for remote and non-invasive genetic priming of solid tumors for synNotch-CAR T treatment. The resulted subpopulation of tumor cells expressing a clinically validated antigen upon Dox-gated FUS stimulation can serve as priming cells and “training centers” for the activation of synNotch CAR T cells, which will then, at the local neighborhood of tumor site, attack the whole population of cancer cells expressing a less specific but widespread tumor antigen, and eradicate the tumor (Fig. 7).

**Figure 7:**
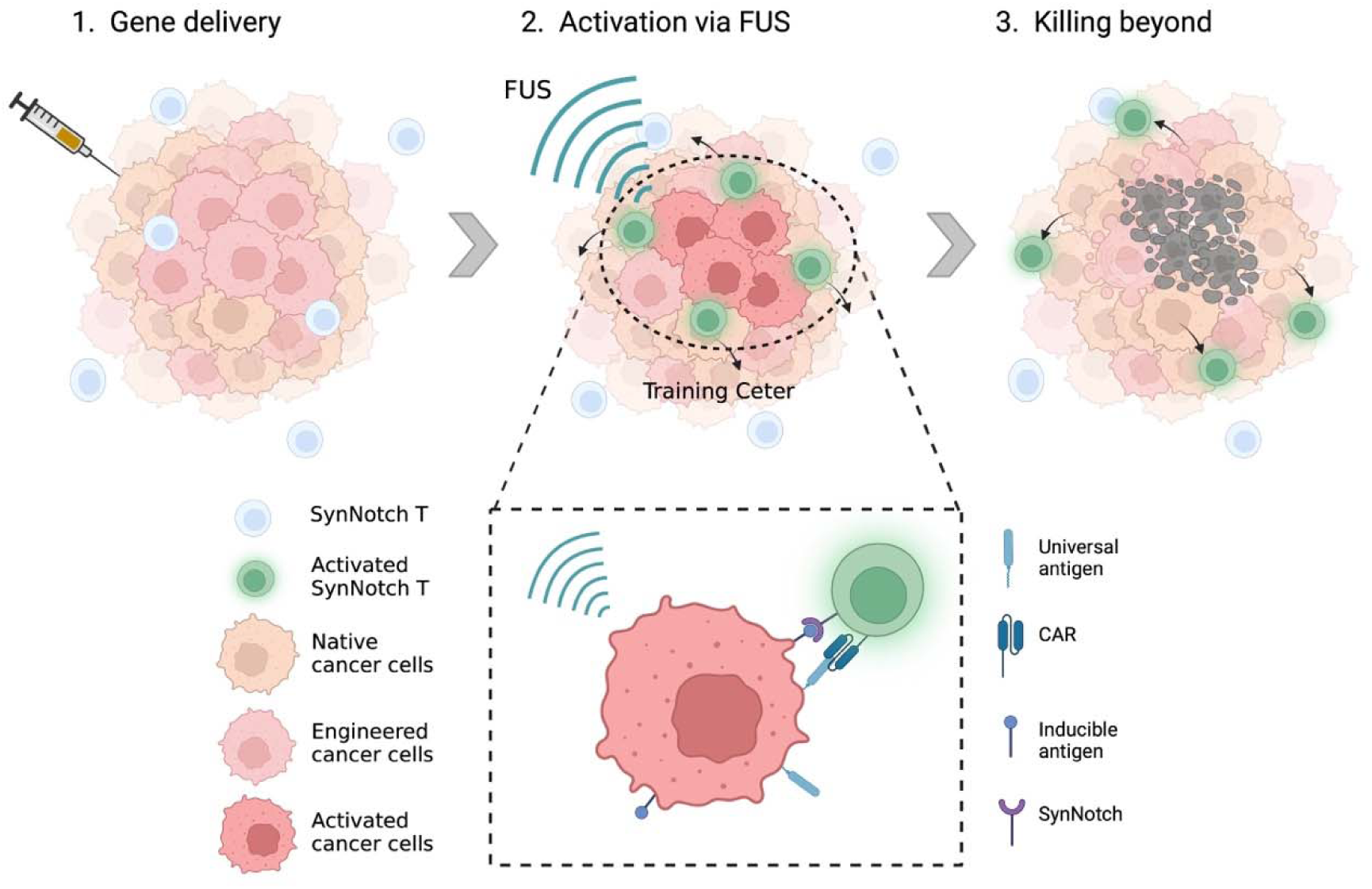
Schematic representation of the combinatorial immunotherapy strategy. This illustration outlines the proposed therapeutic approach, which includes: (1) gene delivery of the CaDox system to vaccinate the tumor; (2) local activation of the CaDox system by focused ultrasound (FUS) stimulation for tumor priming, inducing clinically validated and specific antigen expression on a subset of tumor cells; and (3) the recognition of these inducible antigens by synNotch CAR T cells, leading to the expression of CAR against potent and widespread tumor antigens and the subsequent killing of the whole cancer cell population at the tumor sites.

FUS is a non-invasive technique that generates mechanical energy in the form of ultrasound waves. At the tissue level, FUS can non-invasively open the blood-brain-barrier for targeted drug delivery^52^, induce temporary cellular permeability via sonoporation to enhance drug or gene uptake^53^, and facilitate non-thermal tissue destruction through histotripsy^54^. On a cellular scale, when these waves interact with cells, they can cause mechanical stimulation, which in turn can trigger various mechanotransduction pathways including integrin signaling^55^, the FAK/MAPK/ERK signaling cascade^56, 57^, and the activation of mechanosensitive ion channels like Piezo1^58, 59^, MscL^60, 61^, or TRPA1^21^. The activation of these ion channels by FUS has been explored for its potential to affect cellular responses. However, thus far, the use of ultrasound to stimulate cells has been primarily limited to modulating endogenous gene expression and neuronal activities, which typically lacks the specificity of genetic and cellular controls. To the best of our knowledge, this study is the first to demonstrate the feasibility of FUS-induced mechanical stimulation to directly control user-designed gene expression and cellular functions without the need of a cofactor such as microbubbles. Our study hence represents a new advancement in the field of mechanogenetics and sonogenetics, by demonstrating the versatility and adaptability of FUS-based mechanogenetics, paving the way for more diverse and targeted applications of ultrasound in specific gene regulation and cellular control.

The present system relies on the inherent mechanosensitivity of cancer cells, specifically pannexin1 (PANX1)-mediated calcium responses in prostate cancer PC-3 cells^62^. The activation of PANX1 leads to ATP release, which in turn stimulates P_2_Y purinergic receptors. This stimulation results in an increase in inositol 1,4,5-triphosphate (IP_3_) levels and the subsequent release of calcium, further activating PANX1 channels and releasing ATP, thereby establishing a positive feedback loop. Our findings are consistent with the existing understanding of the role of PANX1 channels in initiating and propagating intercellular calcium waves (see supplementary Fig. 2). PANX1 can thus facilitate the conversion of localized stimulation by FUS into synchronized response from the whole targeted tissue masses, such as solid tumors. This unique aspect of PANX1 channels, as a sensor protein for mechanogenetics, sets them apart from mechanosensitive ion channels that only trigger calcium influx into the intracellular space directly sensing the mechanical perturbation^21, 22^.

In the current study, we used PC-3 cells as our model; however, FUS-mediated calcium responses have been observed in various cancer cell types, including breast and bladder cancer cells^32, 33^. The modular design of our FUS-CaDox system can hence be readily extendable for tissue priming of other cancer types. Indeed, the functionality of CaDox system can be observed in different cell types tested, including Jurkat T cells and MDA-MB-231 human breast cancer cells, highlighting its broad applicability (supplementary Fig. 5). Some studies have suggested potential roles for PANX1 in cancer progression; for example, metastatic cell survival during microvascular deformation is facilitated by increased ATP release from mechanosensitive PANX1 channels activated by membrane stretch, which then acts as an autocrine suppressor of deformation-induced apoptosis through P_2_Y-purinergic receptors^63^. Although further investigation is required to gain a mechanistic understanding of the relationship between FUS sensitivity and cancer cell properties, the observation that cancer cells tend to respond to FUS more than normal healthy cells suggests a potentially additional therapeutic advantage for this technique, differentiating cancer cells from those normal ones.

To harness the FUS-mediated calcium response for customized transcriptional activities, we developed an AND logic gate circuit termed CaDox, which relies on both calcium and doxycycline for its activation. CaDox is designed to minimize crosstalk between the synthetic circuit and endogenous signaling pathways by employing a truncated NFAT motif and the Tet-On component. While doxycycline caused some minor genetic leakages in the absence of stimulated calcium *in vitro*, the CaDox circuit is highly effective and only responds significantly in the presence of both FUS-induced calcium and doxycycline (Figs. 2G-J). In our *in vivo* killing assay, the doxycycline-only group exhibited no tumor suppression, while the doxycycline and FUS combined group demonstrated significant suppression (Fig. 6C). This suggests that the minor leakage of Doxycycline of CaDox may not affect practical applications *in vivo* even without extensive optimization effort. If needed, the system’s expression level can be further controlled by doxycycline concentration and duration to mitigate background leakages (Figs. 2G-H), offering flexibility for optimization based on specific application requirements.

We explored the potential of our mechanogenetic system for immunotherapy by rewiring the circuit to express tCD19 antigen upon FUS activation for tumor priming, enabling precise targeting by synNotch CAR T cells. This approach offers a clear advantage over conventional cytotoxic/suicide gene therapy for cancer treatment, where cytotoxic/suicide genes are directly delivered into tumors and hence significantly limited by delivery efficiency^64^. In contrast, our combinatorial immunotherapy strategy utilizes a subset of infected and induced cancer cells to serve as “training centers”, which activate synNotch CAR T cells for the recognition and eradication of the whole cancer cell population at the tumor region. Furthermore, the CAR expression on the synNotch CAR T cells activated by the local “training centers” at the tumor site is transient in nature and will hence gradually decay when the synNotch CAR T cells stride away from the priming site^49^. This will hence allow the use of highly potent but less specific CAR without the off-tumor toxicity yet achieving a high treatment efficiency. Although the use of tCD19 as an initiator for synNotch CAR T cells might cause non-selective killing of B cells expressing CD19 (i.e. B cell aplasia), lymphodepletion—a well-established process in CAR T-cell therapy—minimizes the number of B cells in the patient’s body to create a time window for treatment^65^. In fact, in clinical trials, fludarabine and cyclophosphamide have been established to pretreat patients for lymphodepletion before CAR T application^66^. B cell aplasia itself is also clinically manageable with immunoglobulin replacement^67, 68^. Given the rapid advancements in the field of immunotherapy and the potential impact of our mechanogenetic system, there is a promising horizon for further refinement and integration of this approach into a broader array of clinical applications, paving the way for safer and more effective cancer treatments in the future.

## Methods

### Animals

Animal experiments were conducted in accordance with Protocol S15285, approved by the UCSD Institutional Animal Care and Use Committee (IACUC). All researchers involved in animal works adhered to relevant animal-use guidelines and ethical regulations throughout this study. NOD scid gamma (NSG) mice (6–8 weeks old, male), obtained from UCSD Animal Care Program (ACP), were utilized for all animal experiments.

### Cloning

Plasmids were generated using either the Gibson Assembly (New England Biolabs, NEB, E2611L) method or the T4 DNA ligation (NEB, M0202L) method, depending on the specific requirements of each construct. Throughout the study, the appropriate cloning strategy was employed for each construct, and the resulting plasmids were used for subsequent experiments as described in the relevant sections of the Methods and Materials. All plasmids utilized in this study are detailed in Supplementary Table 1.

### Site-directed mutagenesis of NFAT for variant calcineurin binding sequences

To elucidate the kinetic differences associated with varying affinities to calcineurin (CN), we engineered several NFAT variants, each harboring distinct CN binding sequences. These modifications were introduced using the Q5 Site-Directed Mutagenesis Kit (NEB, E0554S), following the manufacturer’s instructions. The primers for the mutagenesis processes were designed via the NEBaseChanger website (https://nebasechanger.neb.com), ensuring both specificity and efficiency. Following mutagenesis, the derived NFAT variants underwent Sanger sequencing for verification, confirming the intended modifications in the CN binding sequences.

### Cell culture and Reagents

The cell lines used in this study, including HEK293T, PC-3, Jurkat T, and MDA-MB-231 cells, were obtained from the American Type Culture Collection (ATCC). HEK293T, PC-3, and MDA-MB-231 cells were cultured in Dulbecco’s Modified Eagle Medium (DMEM, Gibco, 10569069) supplemented with 10% heat-inactivated FBS (HI-FBS, Gibco, 16140071) and 1% penicillin/streptomycin (P/S, Gibco, 15140122). Jurkat T cells were maintained in Roswell Park Memorial Institute Medium (RPMI-1640, Gibco, 22400105) supplemented with 10% HI-FBS and 1% P/S. Primary T cells were cultured in X-Vivo™ 15 medium (Lonza, BE02-060Q) supplemented with 5% HI-FBS, 50 μM 2-Mercaptoethanol (Gibco, 31350010), and 1% recombinant human IL-2 (Peprotech, 200-02). All cells were maintained in a humidified incubator at 37°C with 5% CO2 and passaged as per standard protocols to ensure optimal growth and experimental conditions.

### Exogenous gene delivery and establishment of stable cell lines

To investigate the kinetics of NFAT variants in HEK293T cells, we used Lipofectamine 3000 kit (Thermo Fisher, L3000015) to transiently deliver exogenous genes into cells, following the manufacturer’s guidelines. Briefly, we mixed the Lipofectamine reagent with DNA plasmids encoding the exogenous genes of interest and incubated the mixture at room temperature for 15 minutes to allow for complex formation. We then added the complex to the cells and incubated them for 24-48 hours before performing timelapse imaging.

To engineer cell lines for stable expression of exogenous genes such as CaDox components, we use viral transduction based on lentiviral methods. Briefly, we produced lentiviral particles by co-transfecting Lenti-X packaging cells with the transfer vector encoding the gene of interest, along with packaging plasmids (pVSV-G and pΔR) using the calcium phosphate method (Promega, E1200). After 48-72 hours, the supernatant containing the lentiviral particles was collected, filtered, and concentrated using the Lenti-X Concentrator (Takara, 631232). The resulting viral pellet was resuspended in 100 µL of PBS and stored at −80°C until further use. For transduction, target cells were seeded in a 6-well plate and grown to 70-80% confluency. The cells were then transduced with the concentrated lentiviral particles. Transduced cells were subsequently selected using FACS sorting (Sony, SH800) to establish a stably expressing cell population.

### Primary T cell isolation and transduction

Peripheral blood mononuclear cells (PBMCs) were obtained from buffy coats provided by the San Diego Blood Bank and isolated using a lymphocyte separation medium (Corning, 25-072-CV), following the manufacturer’s guidelines. Primary human T cells were then extracted from the PBMCs using the Pan T Cell Isolation Kit (Miltenyi, 130-096-535) and activated with Dynabeads Human T-Expander CD3/CD28 (Gibco, 11141D) on the same day. For transduction, concentrated lentivirus was added to the T cells on day 3 at a multiplicity of infection of 10 and subjected to spinoculation in a 24-well plate coated with 30 µg/mL RetroNectin® (Takara, T100B). T cells continued to expand, and Dynabeads were removed on day 6. When required, transduced T cells were sorted using a fluorescence-activated cell sorting (FACS) machine (Sony, SH800) to acquire a pure population.

### Fabrication and characterization of ultrasonic transducers

For the *in vitro* study, we fabricated two ultrasound transducers in-house using standard methods described previously^69^. The first was a lithium niobate (LiNbO_3_) pressed-focused 35-MHz transducer (f-number = 1, focal length = 6 mm). Higher frequency transducers offer advantages for mechanistic studies. A higher frequency results in a narrower focused area, which allows precisely localized stimulation. Moreover, as the mechanical response of cells to ultrasound is often more pronounced at higher frequencies, less power is required to induce cellular responses. This is related to the frequency-dependent nature of ultrasound interaction with biological tissues, where higher frequencies exhibit greater acoustic attenuation^15^.

We also designed and fabricated a highly focused single-element 2-MHz ultrasound transducer (f-number = 0.67, focal length = 6 mm). This transducer was utilized to examine cellular responses at a frequency that is clinically relevant, providing important insights for potential therapeutic applications. Modified hard PZT (Del Piezo, DL-47) was used for this transducer, which is known for its high-power handling capability.

For animal study, we constructed a focused 1-MHz single element transducer in-house using a pre-focused modified PZT (diameter: 70mm, a radius of curvature: 65mm, DL-47, Del Piezo) with a 20 mm hole in the center based on the design from a previous report^70^.

Transducer characterization was conducted by measuring axial and lateral resolution, as well as intensities per applied voltage, using a needle hydrophone (Onda, HGL-0085) (Fig. 4, Supplementary Fig. 1 and 7). The peak negative pressures (PNP) and spatial-peak, temporal-average intensity (I_SPTA_) served as indicators of the strength of the focused ultrasound used in each experiment.

### *In vitro* ultrasound stimulation

For *in vitro* studies, we used a FUS stimulation setup integrated with an inverted epifluorescence microscope. We utilized press-focused 35-MHz and 2-MHz FUS single-element transducers, which were manipulated by a 3D linear stage controller (Sigma Koki, SHOT-304GS) for precise position control. User-defined input signals were generated using a function generator (Stanford Research Systems, SG380) and were amplified by a 50dB power amplifier (E&I, 325LA) before being fed into the ultrasonic transducers. To position the cells at the transducer’s axial focus, we employed a pulse-echo method using a pulser-receiver (Olympus, 5072PR) and oscilloscope (Teledyne Lecroy, 610Zi) to determine the distance between the transducer and culture dish that yielded the maximum echo signal. To align the transducer’s lateral focus with the center of the CCD camera of the microscope, we utilized an acoustic trapping method described previously^71^.

### *In vivo* ultrasound apparatus and stimulation

We developed an in-house FUS stimulation system for animal studies using a 1-MHz transducer to induce localized mechanical perturbation. A coupling cone (length: 65mm) with a 4mm diameter opening at the tip was designed and 3D-printed to hold degassed water along the acoustic path to the animals (See Supplementary Fig. 7). The opening at the end was sealed with an acoustically transparent Mylar film (thickness: 2.5 µm, Chemplex, 100) to prevent water leakage. Custom Python code was written to control a function generator (Stanford Research System, SG386) by sending user-defined input parameters and repetition patterns to the generator. The signal was then amplified by a 50dB power amplifier (E&I, 325LA) before being fed to the transducer. A set of manual translational stages were employed to control the transducer’s focus. Additionally, a feedback-controlled temperature pad (Auber Instruments, WSD-30B) was used to warm the animal bed to maintain the animal’s body temperature under anesthesia. An acoustic absorber (Precision Acoustics, Aptflex F28) was placed under the animal to reduce any reflections.

For the stimulation procedure, doxycycline (1 mg/kg) was first injected into mice 5 to 7 minutes before anesthesia to allow sufficient circulation in the body. The anesthetized mouse was then placed on the animal bed with its left side on top, and acoustic gel (Aquasonic, 26354) was applied to the tumor area as a coupling media that fills potential air gaps to maximize the transmission of FUS. If necessary, the region was shaved using a handheld trimmer before gel application. The 1-MHz transducer was positioned over the marked tumor area with the tip of the coupling cone placed within 0.5mm of the skin.

For real-time thermometry, a needle-type thermocouple (29 ga., time constant: 0.25 sec, MT-29/2HT, Physitemp Instruments) with a thermometer (HH806AU, Omega) was used. During the procedure, the tip of the needle-type thermocouple was inserted subcutaneously from the side into the mouse near the target region, while the focused transducer was placed on top of the tumor to stimulate the area.

### Live-cell calcium imaging and quantification

Live-cell calcium imaging was performed to monitor intracellular calcium dynamics in response to FUS stimulation. Cells were loaded with the calcium-sensitive fluorescent dye, Fluo-4 AM (Thermo Fisher Scientific, F14201), following the manufacturer’s instructions. Briefly, cells were incubated with 5 µM Fluo-4 AM and 0.02% Pluronic F-127 (Thermo Fisher Scientific, P3000MP) in a buffer solution for 30 minutes at 37°C. After incubation, cells were washed in fresh buffer solution to remove excess dye. The loaded cells were then placed on an inverted epifluorescence microscope (Nikon, Eclipse Ti) for imaging. Alternatively, cells were stably transduced with a genetically encoded calcium sensor, R-Geco1 ^72^, to monitor intracellular calcium dynamics in organoids. R-GECO1 was introduced into cells using lentiviral infection, followed by FACS sorting to generate stable cell lines expressing the calcium sensor. Once stable cell lines were established, they were used to generate organoids. Time-lapse imaging was performed to capture the temporal changes in intracellular calcium levels, and the acquired data were analyzed either using ImageJ software (National Institutes of Health) or using CellProfiler ^73^ with custom-written pipelines to segment cells and calculate normalized changes in fluorescence (ΔF/F).

### *In vitro* luciferase-based assay

For the luciferase-based assay, the Dual-Glo^®^ Luciferase Assay System (Promega, E2920) was used to measure the activity of luciferase in the cell samples according to the manufacturer’s instructions. Briefly, engineered HEK293T cells (0.05M) were seeded in a 96-well plate and allowed to adhere overnight. The next day, the cells were treated with various experimental conditions. At the end of the treatment period, the cells were washed with PBS 3 times and replenished with media for incubation before the assay. After 6 hours, the Dual-Glo Luciferase Reagent was added to each well and incubated at room temperature for 3 minutes to lyse the cells and stabilize the luminescent signal. The luminescence from firefly luciferase (Fluc) was measured using a plate reader (Tecan, Infinite M200 Pro). Afterward, the Dual-Glo Stop & Glo Reagent was added to each well, and the plate was incubated for another 3 minutes at room temperature to quench the Fluc signal and generate luminescence from Renilla luciferase (Rluc). The Rluc luminescence was then measured using the same plate reader. The Rluc (constitutive) luminescence values were used to normalize the Fluc (inducible) data, accounting for any variability or cell number, providing a more accurate representation of the experimental effects on luciferase activity.

For the extracellular ATP measurement, we used the CellTiter-Glo® 2.0 cell viability assay (Promega, G9241) according to the manufacturer’s instructions. PC-3 cells on imaging dishes were stimulated with FUS for 30 minutes, applying a pattern of 1 minute every 5 minutes. After stimulation, the culture media were mixed with an equal volume of CellTiter-Glo® 2.0 reagent and incubated at room temperature for 3 minutes to stabilize the luminescent signal. The luminescence was subsequently measured using a plate reader. The data obtained from this assay offered insights into extracellular ATP levels in the presence and absence of FUS stimulation.

### *In vitro* fluorescence-based assay

For the fluorescence-based assay, mCherry expression was used as a readout to assess the inducibility of the CaDox system against various treatment conditions. Engineered cancer cells expressing the mCherry gene under the control of the CaDox system were seeded in imaging dishes and allowed to adhere overnight. The following day, the cells were exposed to different experimental conditions to test the CaDox system’s response. After the treatment period, the cells were washed three times with PBS and replenished with fresh media before incubation. After 6 hours of incubation, the mCherry fluorescence intensity was measured using an inverted epi-fluorescence microscope with a 20x objective. Images were captured and then analyzed using a custom Python code to segment the cells and calculate the sum of intensities from the segmented regions. This approach allowed for the quantification of mCherry expression levels, providing a reliable measure of the CaDox system’s inducibility under the various treatment conditions.

### Organoid generation

Prostate cancer, PC-3, cell line-derived organoids were established and maintained as described previously ^39^. To create the organoids, 50,000 PC-3 cells were mixed with 10 µL advanced DMEM/F12 media (Life Technologies, 12634010) supplemented with 10 mM HEPES (Life Technologies, 1560106), 2 mM Glutamax, and 1% P/S. The cell mixture was added to 40 µL of Matrigel HC (Corning, CB354248) on ice and then added to the center of a glass-bottom dish. After 20 minutes of incubation at room temperature, 2 mL of complete organoid media (see Supplementary table 2) containing 10 µL Y-27632 dihydrochloride was added to the dish. The dish was then kept in a CO2 cell culture incubator with the dish upside down. The media was replenished every 3-4 days, and after 5-7 days, the complete organoid media without Y-27632 dihydrochloride was used to maintain the cultures. The experiments were conducted on full-grown organoids (12-15 days).

### Acoustic wave simulation

The simulations were performed using COMSOL Multiphysics 6.0 software and the finite element method (FEM), a numerical technique for solving partial differential equations by discretizing the domain into smaller elements. In our study, the frequency-domain pressure acoustic module within COMSOL was employed to simulate the ultrasound field, as this module is specifically designed to analyze pressure wave propagation and interactions in various media. To ensure the accuracy and reliability of the simulations, material parameters related to ultrasound, such as density, acoustic impedance, and attenuation, as well as animal body properties like skin thickness, normal tissue characteristics, and tumor tissue properties, were sourced from relevant literatures^74–78^. This approach allowed us to create a more realistic representation of the materials and conditions in our simulations, ultimately leading to a better understanding of the acoustic wave propagation within the animal body and the amount of energy deposited at the focus of the ultrasound.

### SYTOX-based cell viability assay

To visualize and assess synNotch T cell-mediated killing of antigen-inducible cancer cells *in vitro*, SYTOX blue dead cell stain (Thermo Fisher, S34857) was utilized. SYTOX blue dead cell stain is a cell-impermeant nucleic acid dye that selectively binds to the DNA of dead cells with compromised cell membranes, enabling the visualization and quantification of cell death in real-time imaging experiments. Engineered PC-3-CaDox-tCD19-PSMA+ cell lines were seeded onto 35 mm culture dishes 24 hours before stimulation. On the following day, the seeded cells were treated with or without doxycycline and then subjected to FUS stimulation at designated locations on the dishes. After removing the drug with DPBS, synNotch T cells were added to the dishes in media containing 25 nM SYTOX. Time-lapse images were captured every 5 minutes for a duration of 24 hours to monitor the viability of the PC-3 cells.

### *In vivo* bioluminescence imaging

*In vivo* bioluminescence imaging (BLI) was conducted using an In vivo Imaging System (IVIS) Lumina LT Series III (PerkinElmer). For nano luciferase (NLuc) imaging, Nano-Glo® In Vivo Substrate (Promega, FFz, CS320501) was injected intraperitoneally at a volume of 100 µL of reconstituted solution (0.44 µmoles), following the manufacturer’s guidelines. Imaging commenced 5 minutes after injection and continued until the peak signal was detected. For Firefly luciferase (Fluc) imaging, D-luciferin (GoldBio, LUCK) was administered intraperitoneally at a dose of 150 mg kg-1. BLI began 10 min post-substrate injection and continued until the maximum signal was reached. Living Image software (PerkinElmer) was employed for image analysis. The integrated Fluc luminescence intensity within the tumor region was quantified and subsequently normalized based on the initial measurement of the same tumor, resulting in the normalized tumor size for monitoring tumor growth.

### Software and statistical analysis

Data were graphed and the corresponding statistical analysis was performed in GraphPad Prism 10.0.0. Flow cytometry data were analyzed and plotted using FlowJo 10.8.2. The detailed statistical test methods were indicated in the corresponding figure legends. Schematic figures were created with BioRender.com.

## Data availability

The main data supporting the results of this study are available within the paper and its Supplementary Information. Other raw data generated during this study are available from the corresponding authors on reasonable request.

## Code availability

The code repository for the PID controller and the device interfaces for the in-house built FUS system can be found at https://github.com/phuongho43/ultrasound_pid. The code for the image analysis and quantification can be found at https://github.com/phuongho43/cytomata.

## Acknowledgements

This work was supported in part by grants from NIH EB029122, GM140929, HL121365, HD107206, and CA262815 to Y. Wang. This work was also supported by an NINDS R24 Core Grant and funding from NEI, and NIH/NCI T32 CA009523.

## Author contributions

C.W.Y. and Y.W. conceived and designed the experiments; C.W.Y., C.S., D.N.M.N., L. Z., G.L., A.Z., and A.L. performed the experiments; C.W.Y., C.S., A.L., and P.H. analyzed the data; C.W.Y., L.Z., Z.H., R.C., Y.Z., and N.S. contributed materials; C.J., K.K.S., and Q.Z. provided guidance and feedback. C.W.Y. and Y.W. wrote the paper. All authors reviewed the manuscript and approved the final version.

