## Supplementary Data for "Tumor Priming by Ultrasound Mechanogenetics for with SynNotch CAR T Therapy"

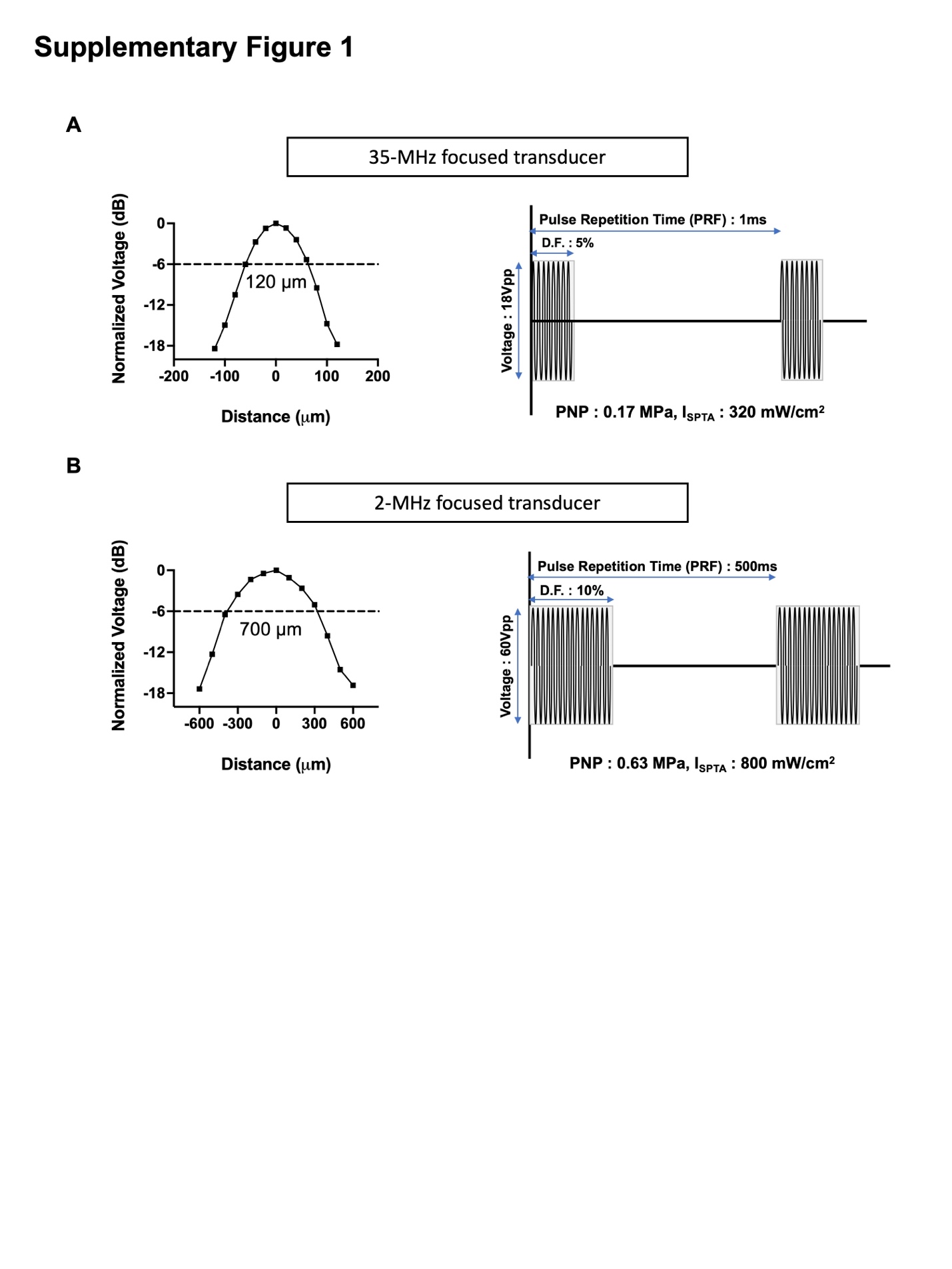

**Supplementary Figure 1: Characterization of FUS transducers for *in vitro* studies.** Beam profile plots at the focal plan, input parameters, pulsed signal patterns, and output powers for (A) 35-MHz single element lithium niobate ultrasound transducer and (B) 2-MHz single element PZT ultrasound transducer. Beam profiles and output powers, represented as peak-negative-pressure (PNP) and spatial-peak-temporal-average intensity (I_SPTA_), were measured using a hydrophone. Note that both transducers were operated within the range of low-intensity pulsed ultrasound scheme.

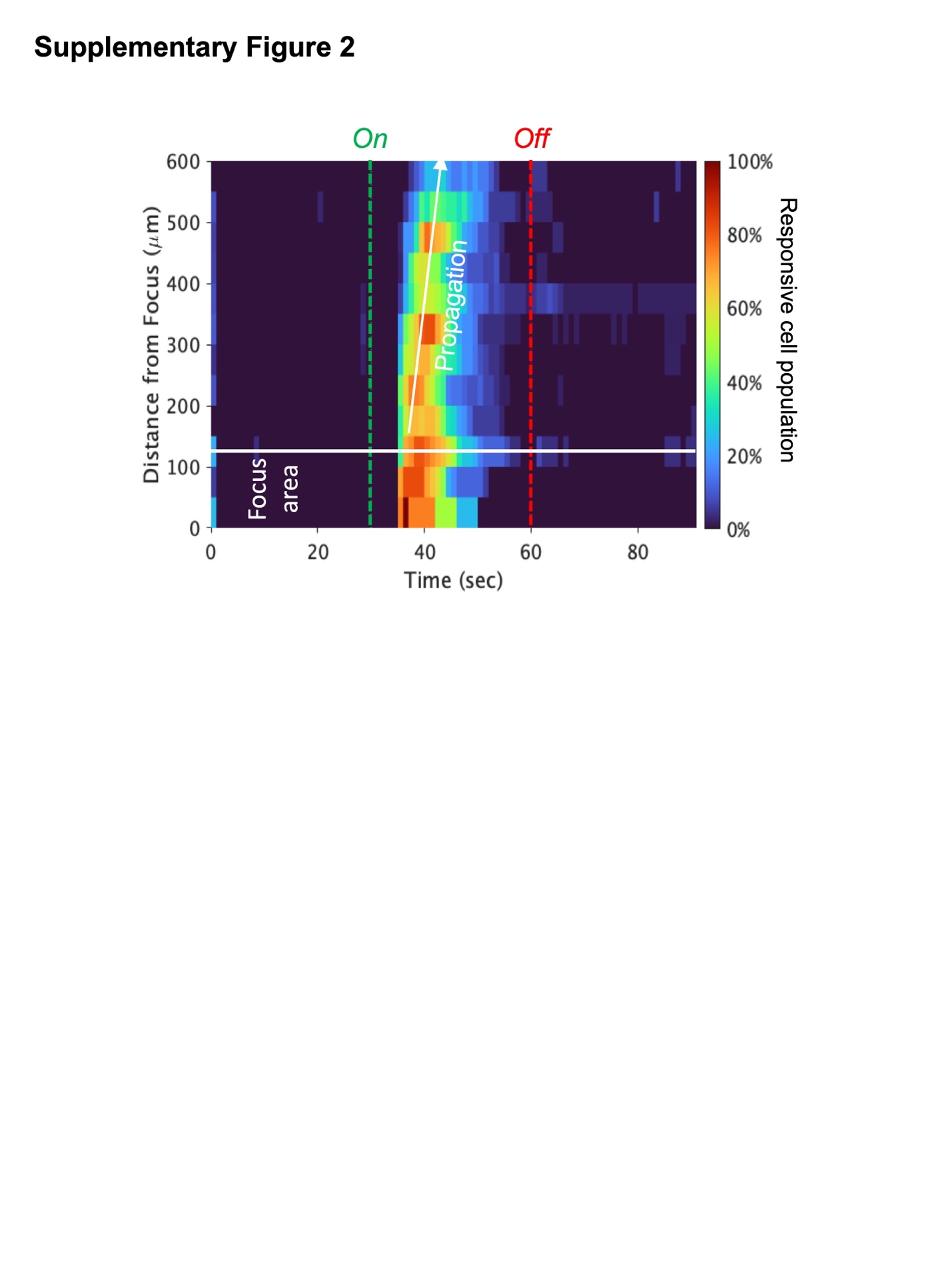

**Supplementary Figure 2: Bivariate histogram of FUS-induced calcium responses.** The histogram presents the proportion of cells that respond over time, with respect to their distance to the focal point of the transducer. Green and red dotted lines indicate the FUS on and off timings, respectively. The responses were detected beyond the FUS focal point and propagated to distances exceeding 500 µm.

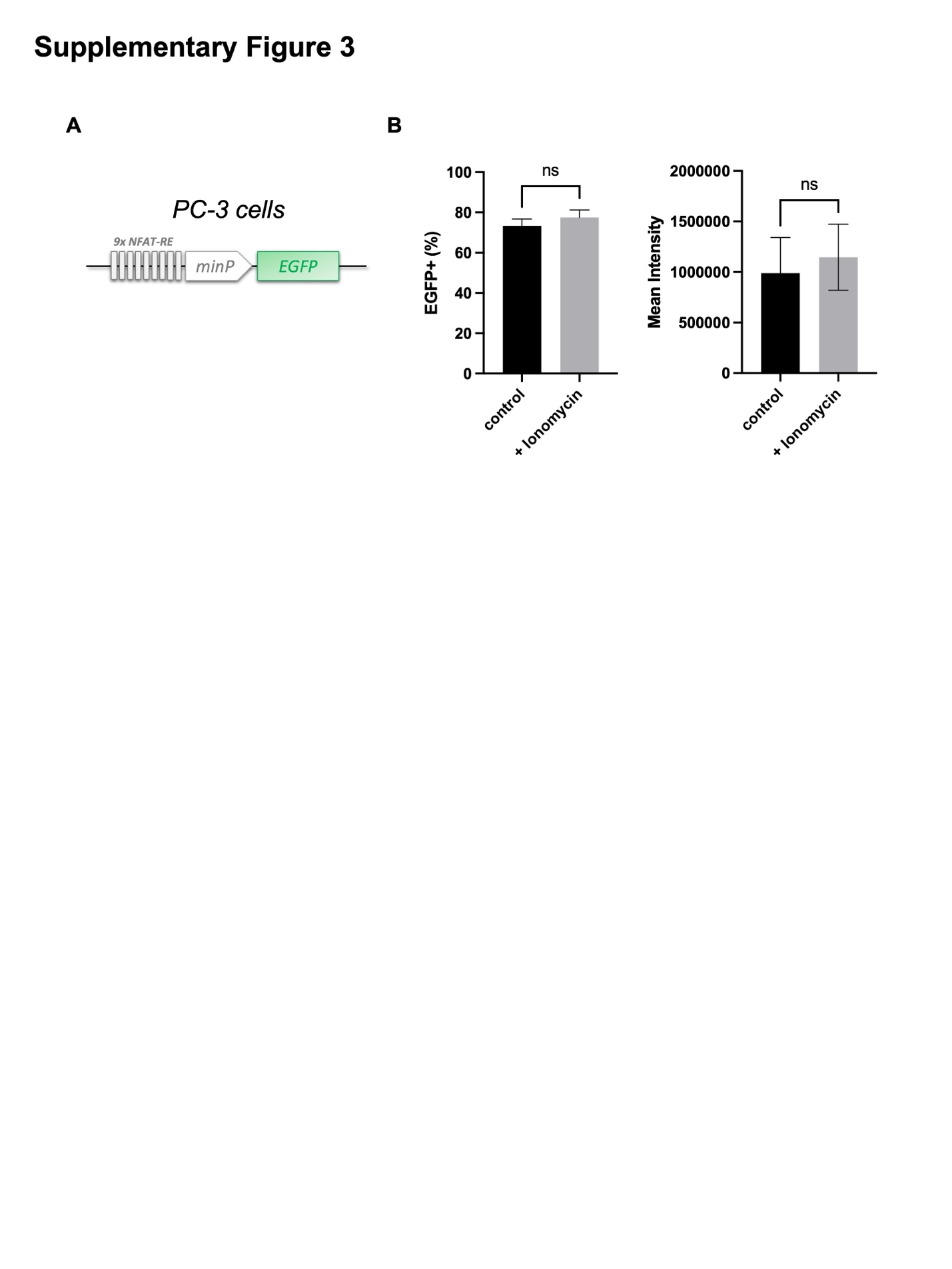

**Supplementary Figure 3: Genetic leakages from the NFAT-RE based circuit.** (A) The circuit features nine repeats of NFAT response elements (NFAT-RE) followed by a minimal promoter, with EGFP as the inducible reporter gene. (B) Characterization of the NFAT-RE-based gene circuit with and without ionomycin (1µM) in PC-3 cells. n=3. Statistical significance was determined using a t-test.

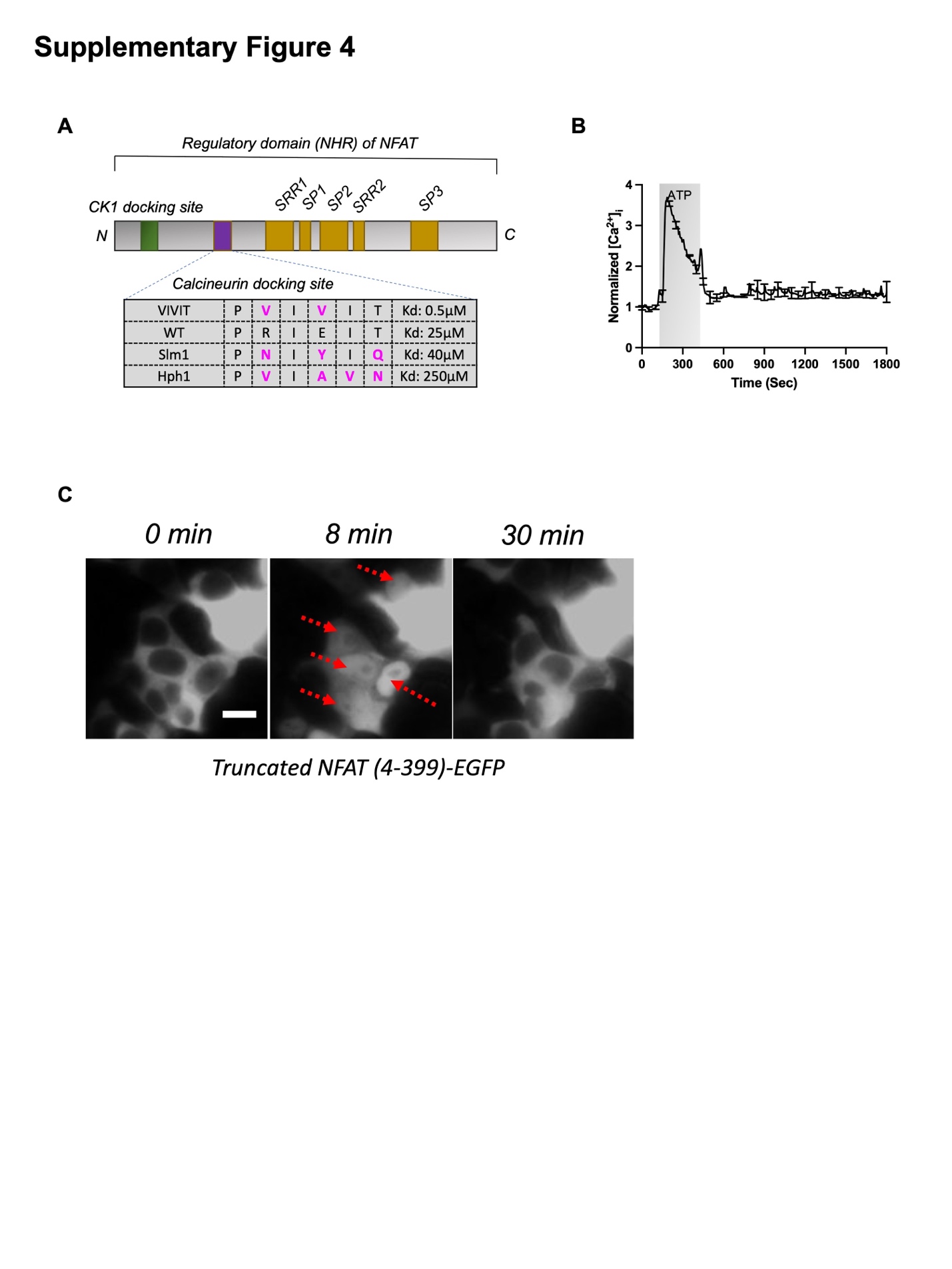

**Supplementary Figure 4: Engineering of NFAT mutants with alterations at the calcineurin docking site.** (A) Schematic diagram depicting the domain structure of the NFAT protein and a list of mutations at the calcineurin docking site. (B) Intracellular calcium response upon incubation with 60 µM ATP. ATP was added at the 2-minute mark and washed off at the 7-minute mark, as indicated by the grey area. (C) Timelapse images of truncated NFAT (4-399)-EGFP upon ATP induction. Red arrows highlight the cells with clear nuclear translocation of truncated NFAT. Error bars represent the standard deviation (SD).

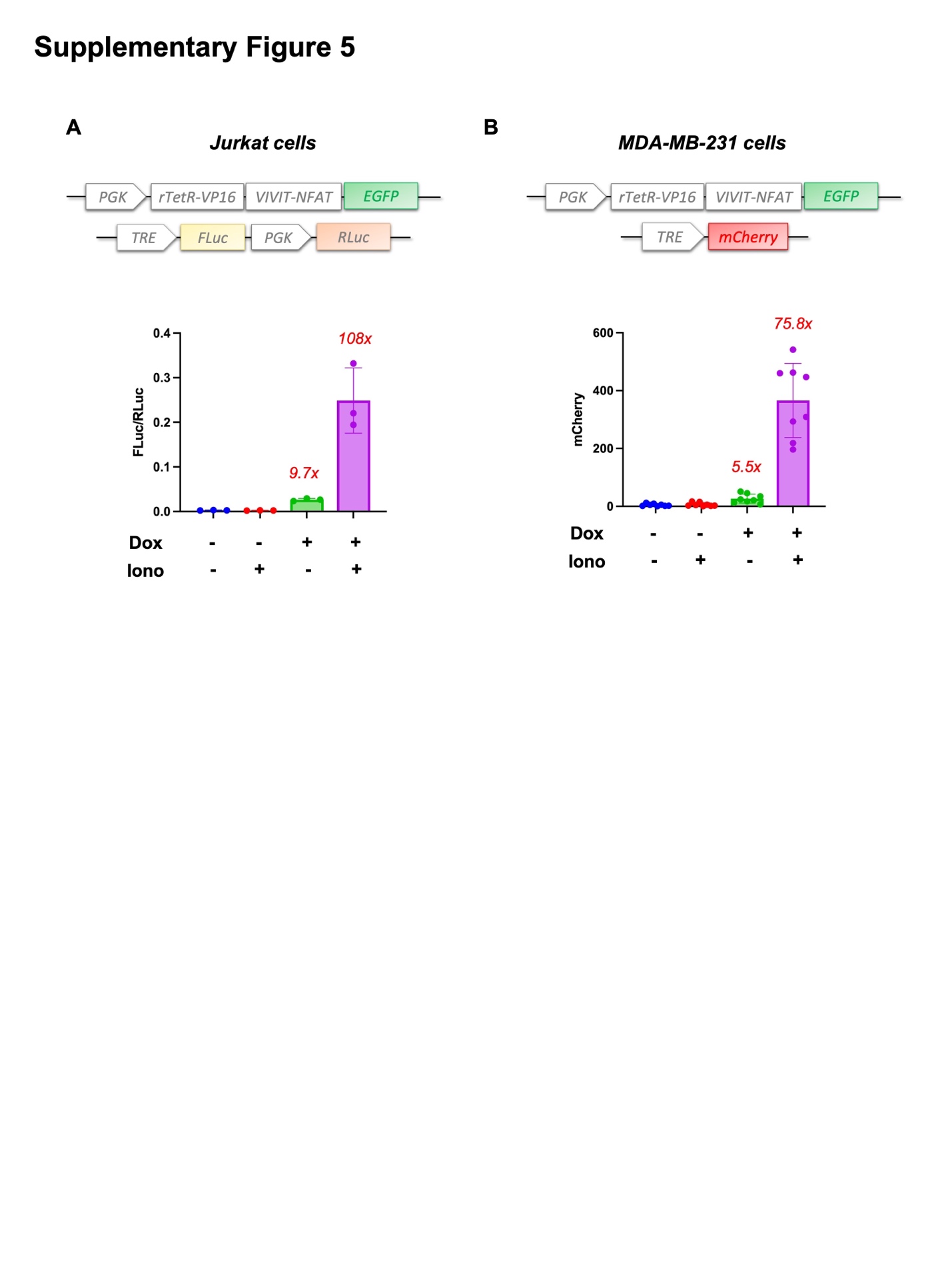

**Supplementary Figure 5: The functionality of CaDox system in different cell types.** The CaDox system demonstrates consistent functionality across various cell types, including (A) Jurkat T cells and (B) MDA-MB-231 human breast cancer cells, highlighting its broad applicability.

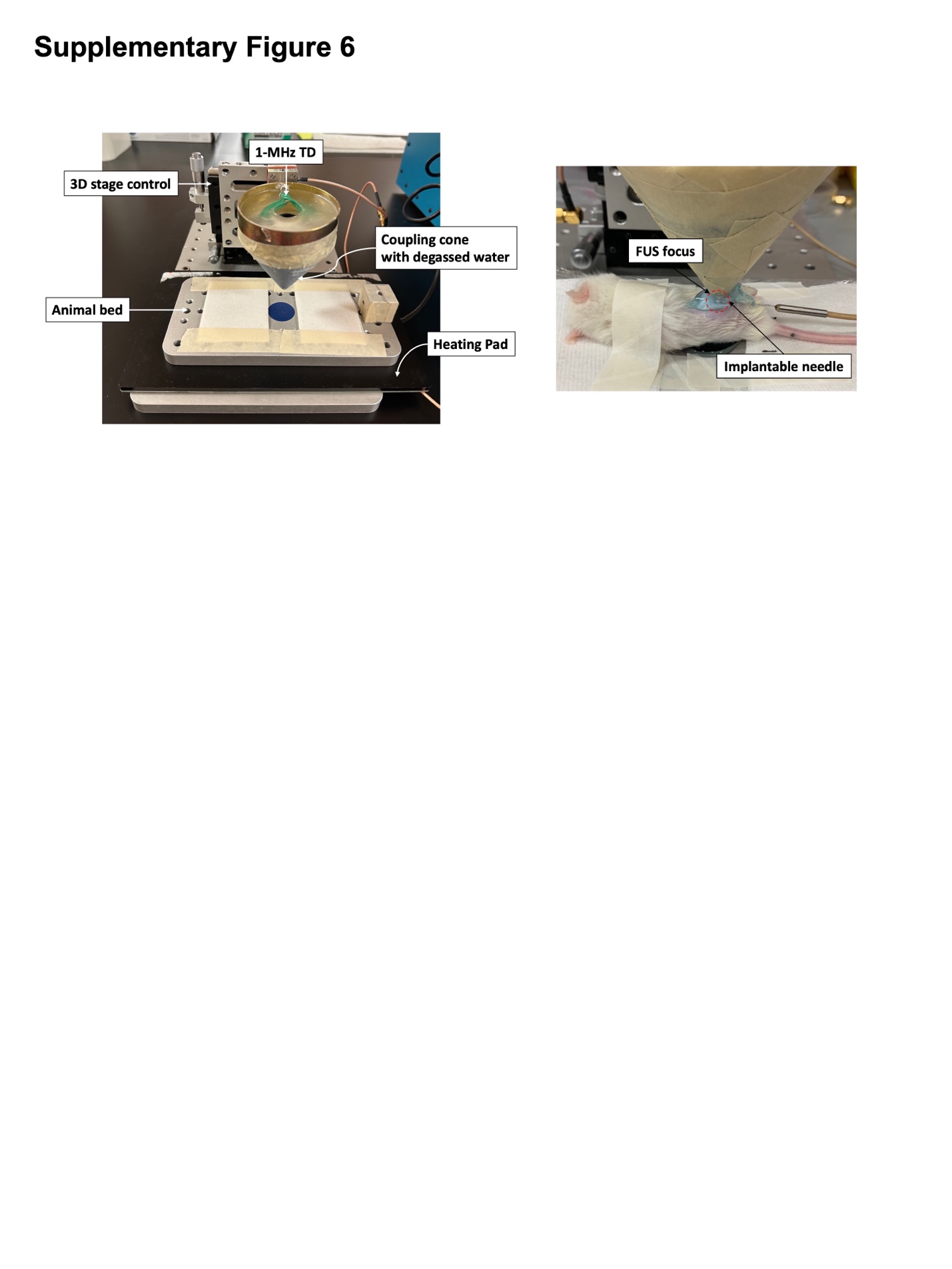

**Supplementary Figure 6: The custom FUS stimulation system for animal studies.** The image displays the in-house developed FUS stimulation system, including the 1-MHz transducer with a 3D-printed coupling cone designed to hold degassed water along the acoustic path to the animals, the feedback-controlled temperature pad used as the animal bed to maintain body temperature under anesthesia, and the set of manual translational stages employed to control the transducer's location. The coupling cone features a 4 mm diameter opening at the tip, sealed with an acoustically transparent Mylar film to prevent water leakage. Additionally, a close-up image of the animal with acoustic gel and the transducer positioned for FUS stimulation is shown at the right side. A needle-type thermocouple was inserted into the animal's body at the FUS focus region to measure the temperature changes induced by the stimulation.

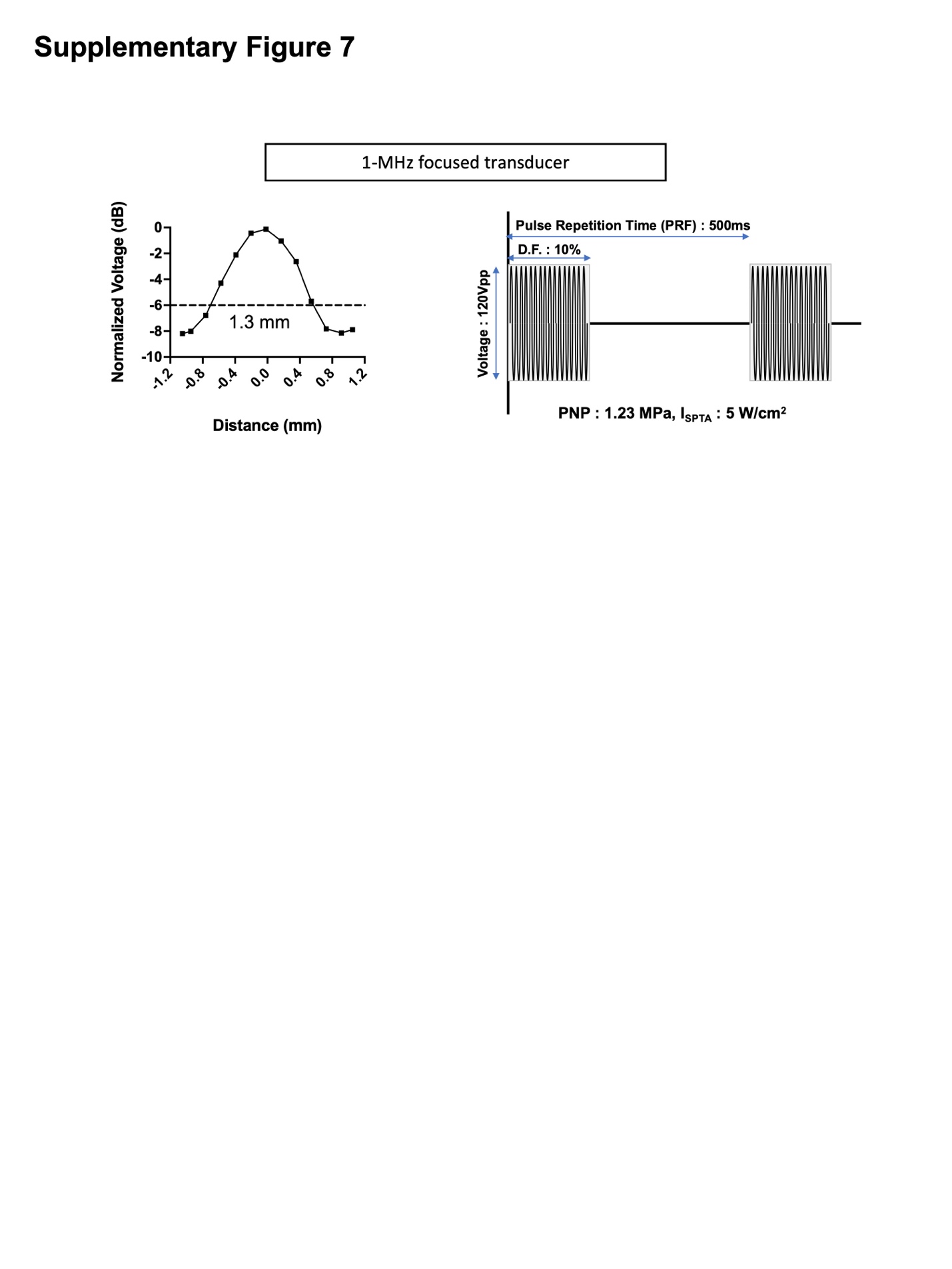

**Supplementary Figure 7: Characterization of FUS transducer for *in vivo* study.** (A) Beam profile plots, input parameters, pulsed signal patterns, and output powers for 1-MHz single element PZT ultrasound transducer.

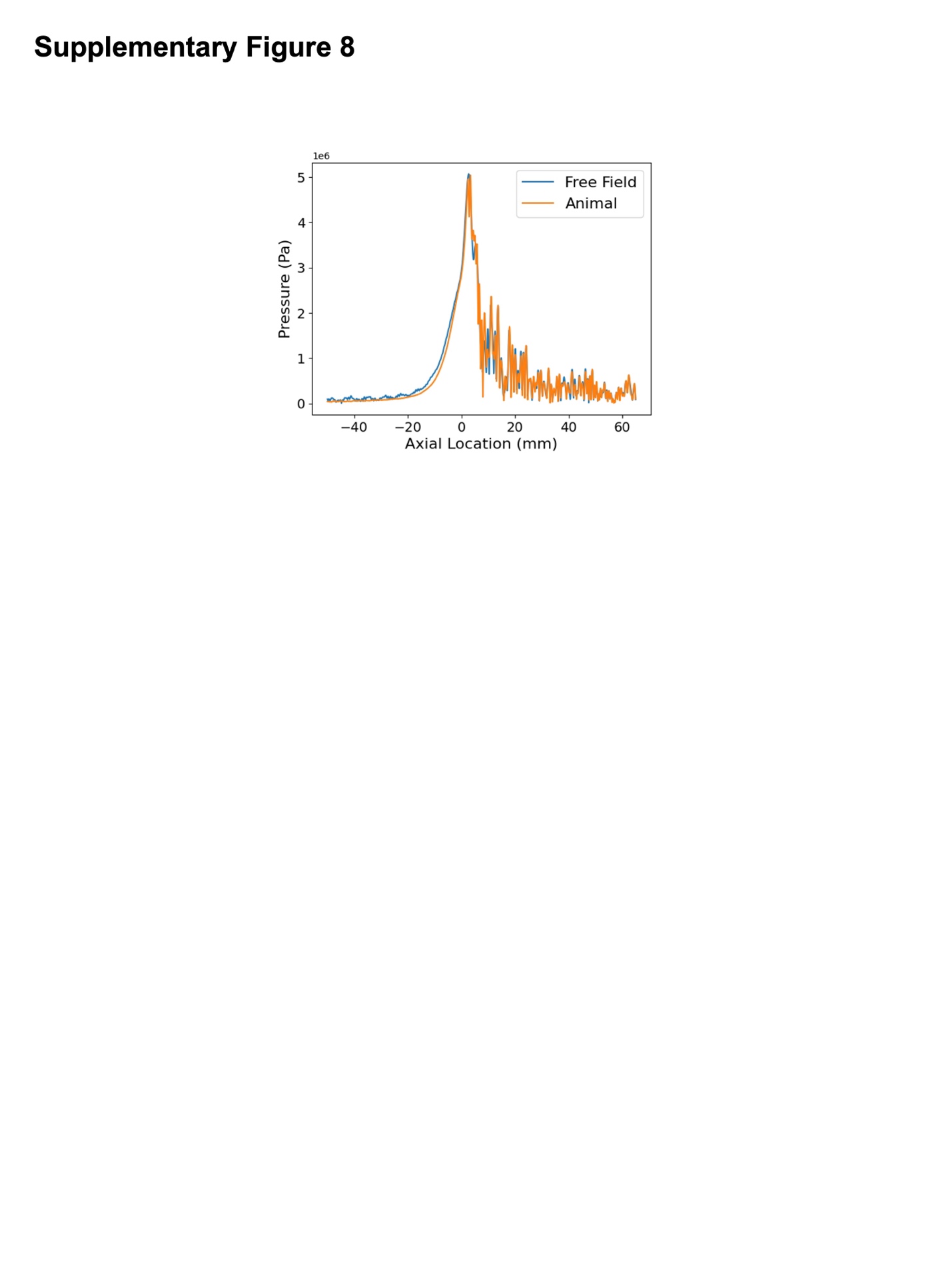

**Supplementary Figure 8:** **Acoustic wave propagation simulation.** Comparison of the pressure field for acoustic wave propagation in two different scenarios: free field (water) and within an animal body consisting of skin, muscle, and tumor. All required parameters for the simulation were referenced from relevant literatures (See Methods). The blue line represents the simulation in the free field, while the orange line illustrates the simulation within the animal body, demonstrating the differences in pressure distribution across the axial direction. The maximum pressure in the free field is approximately 5.071 MPa at a location of 2.71 mm, while the maximum pressure in the animal body is approximately 5.043 MPa at a location of 3.33 mm. The difference in maximum pressure is about -27,653 Pa, with a percentage difference of approximately -0.55%.

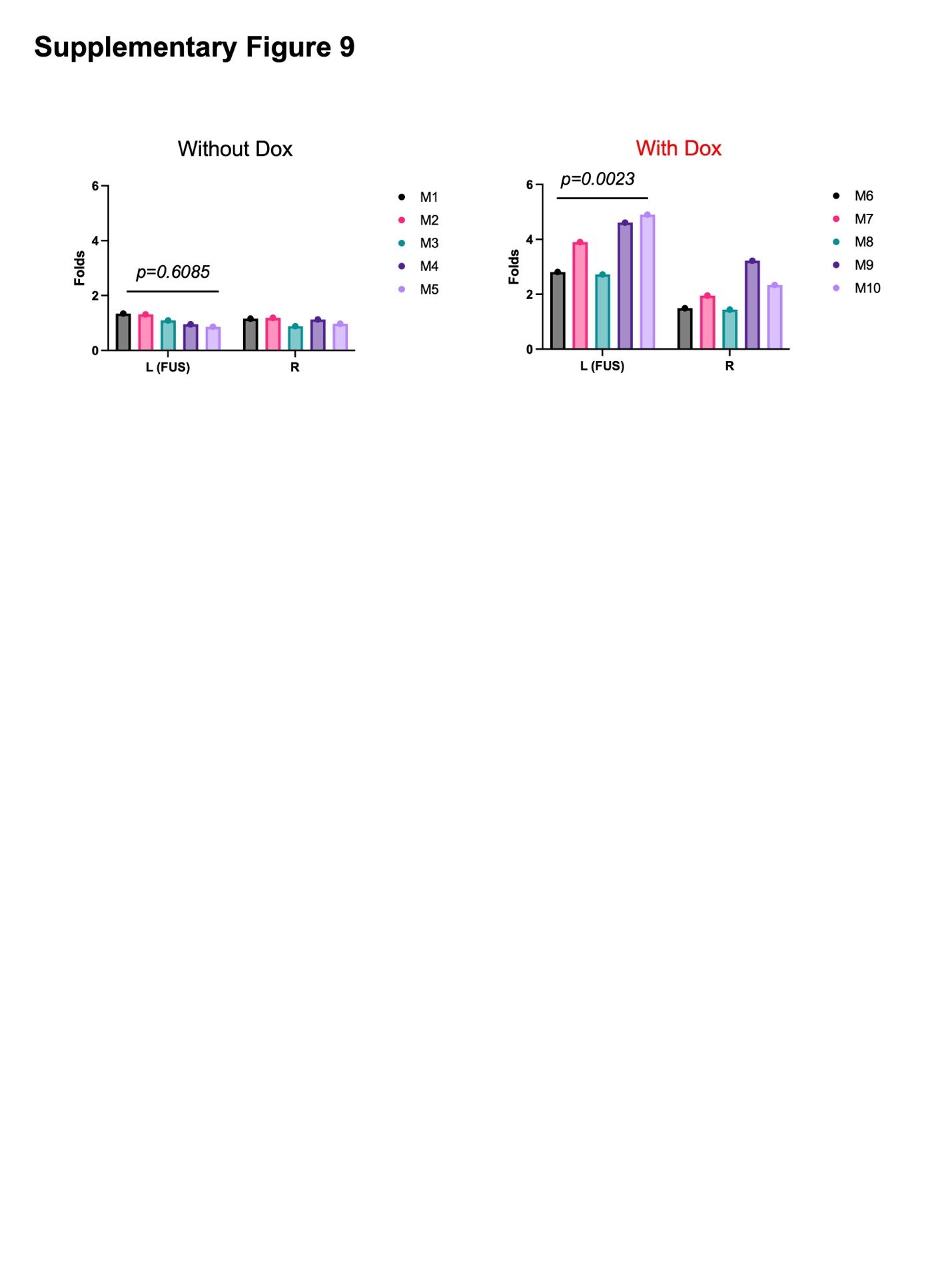

**Supplementary Figure 9: *In vivo* gene expression of the CaDox system in individual mice.** Mice without doxycycline administration (M1 to M5) did not show a difference between tumors with FUS (left tumors) and without FUS (right tumors). In contrast, mice with doxycycline administration (M6 to M10) exhibited a statistically significant contrast between left and right tumors. The gene activations were measured by BLI imaging at 6 hours from the treatment. This indicates that, consistent with *in vitro* assays, mechanical stimulation by FUS can induce gene expression gated by doxycycline.

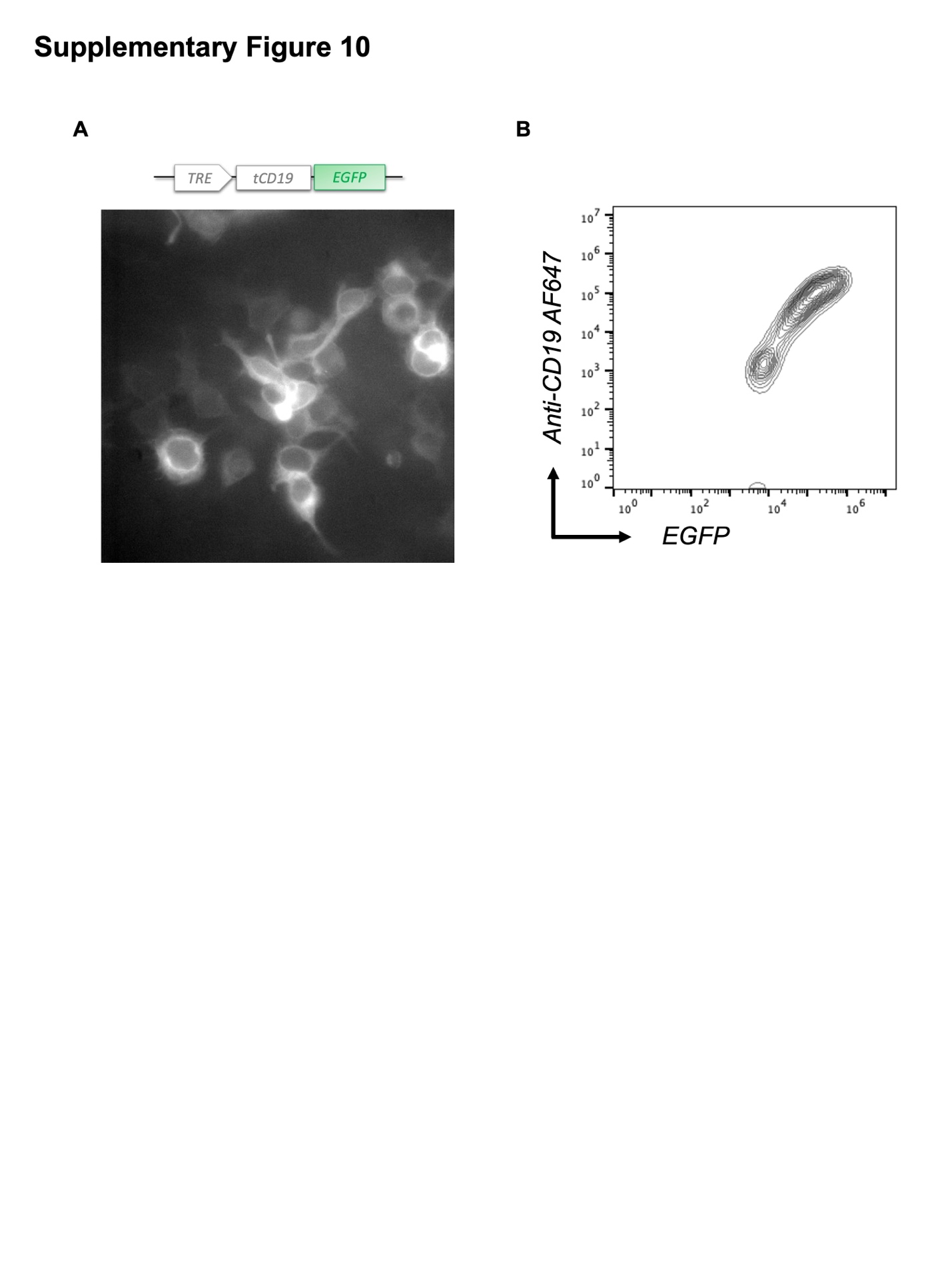

**Supplementary Figure 10: Inducible tCD19 expression in PC-3 cells.** Truncated CD19 was directly fused with EGFP and induced by doxycycline (200 nM) and ATP (60 µM). (A) Representative image of tCD19-EGFP expression on the plasma membrane. (B) Flow cytometry analysis of tCD19 in PC-3 cells, plotting EGFP intensity against AF647 intensity from anti-CD19-AF647 staining. EGFP expression corresponds well with CD19 staining, indicating that the induced tCD19 on the plasma membrane is effectively recognized by the anti-CD19 antibody.

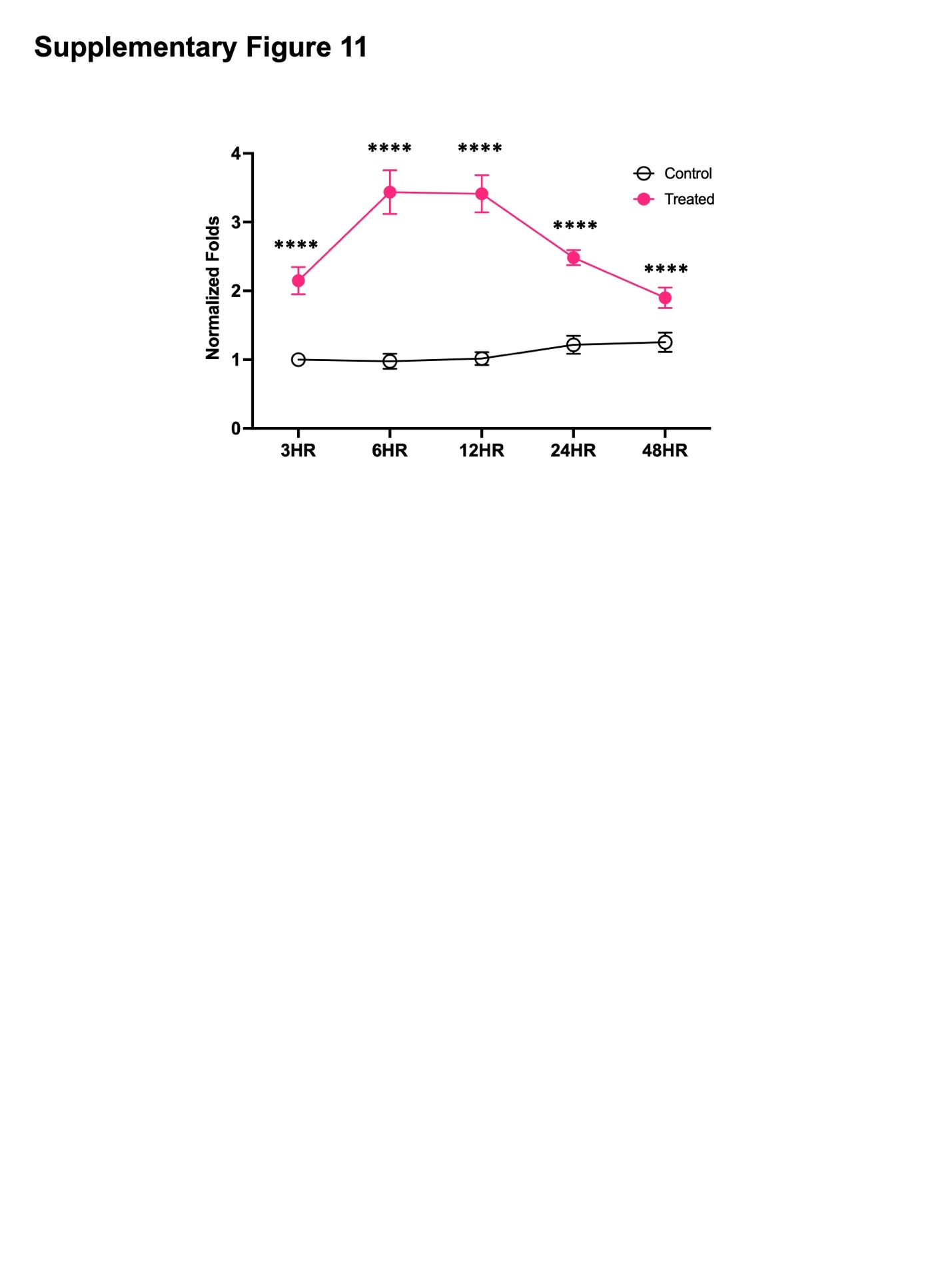

**Supplementary Figure 11: Normalized fold changes of tCD19 expression over 48 hours**. PC-3-CaDox-tCD19-EGFP cells were subjected to various treatments: no treatment (n=10) and combined treatment (n=7). Error bars represent SEM.

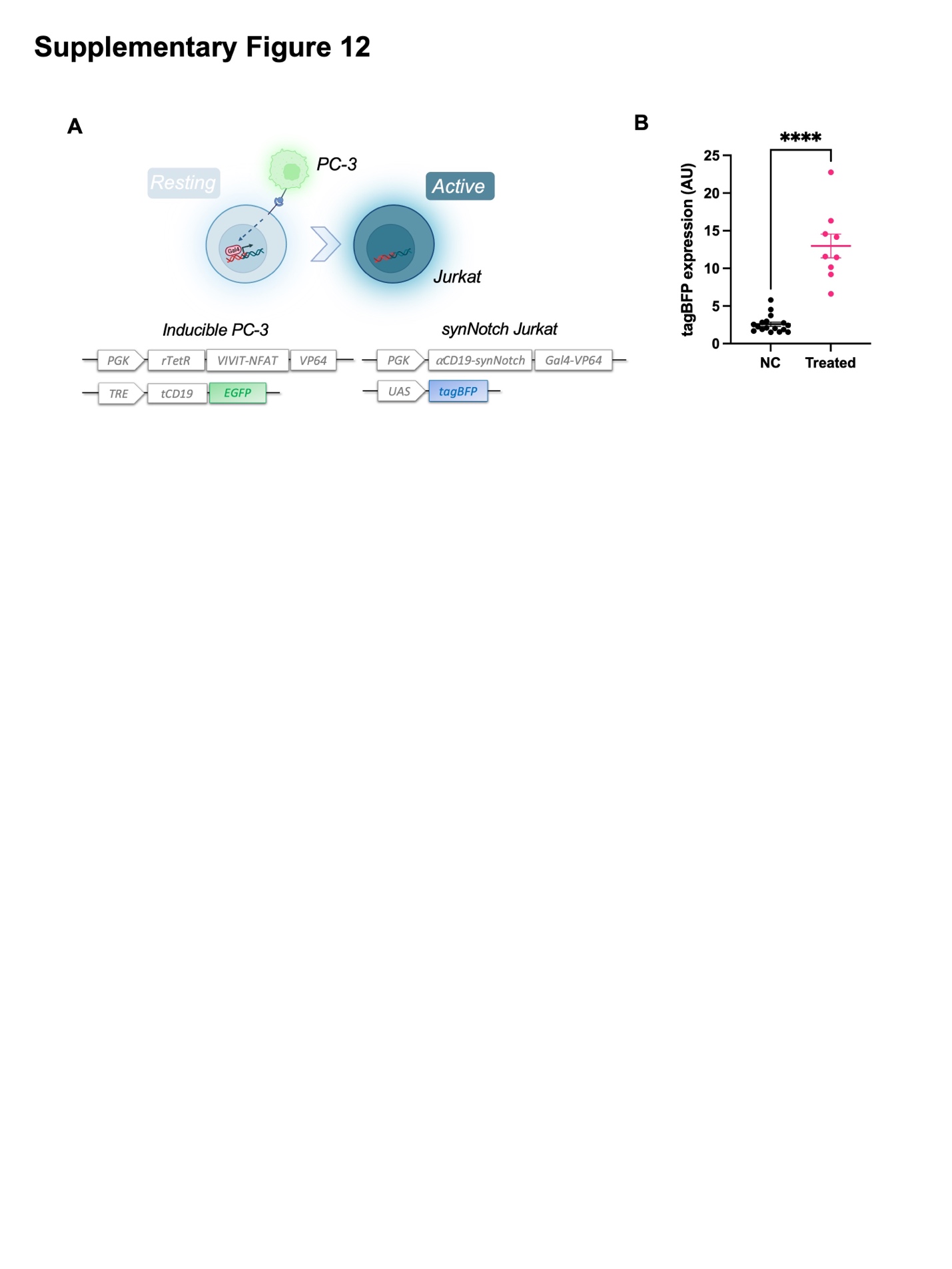

**Supplementary Figure 12: Verification of anti-CD19 synNotch activation against inducible tCD19.** (A) Schematic diagrams of constructs used to engineer synNotch Jurkat T cells for activation by inducible tCD19. (B) Quantification of tagBFP expression in Jurkat T cells activated by tCD19 expressed on PC-3-CaDox-tCD19 cells under different conditions: no treatment (n=17), and combined doxycycline and FUS treatment (n=9). Error bars represent SEM.

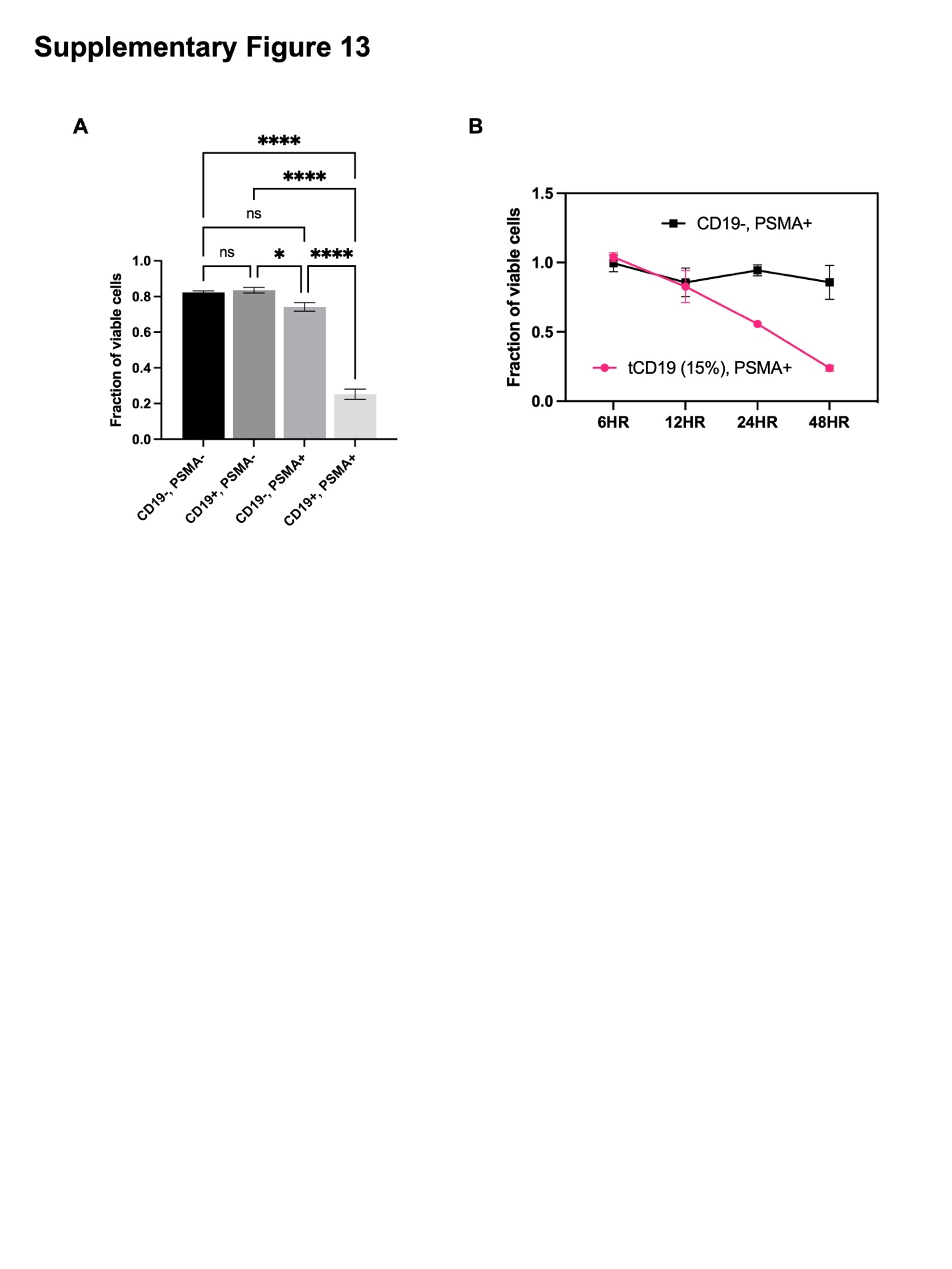

**Supplementary Figure 13: SynNotch CAR T cell cytotoxicity against different control groups.** (A) Viability of various groups of PC-3 cells, CD19^-^/PSMA^-^, CD19^+^/PSMA^-^, CD19^-^/PSMA^+^, and CD19^+^/PSMA^+^ (n=3 for all groups), after co-culturing with synNotch CAR T cells for 24 hours at a 1:1 ratio. (B) Comparison of synNotch CAR-mediated killing of PC-3-PSMA^+^ cells, which show a 15% tCD19 expression from those induced PC-3-CaDox-tCD19 PSMA^+^ cells. This result indicates that a fraction of cells expressing the clinically validated tCD19 can activate synNotch CAR T cells to kill the whole cancer population expressing the common antigen at the tumor site. Error bars represent SEM. Statistical significance was determined by ANOVA with Tukey's multiple comparison test.

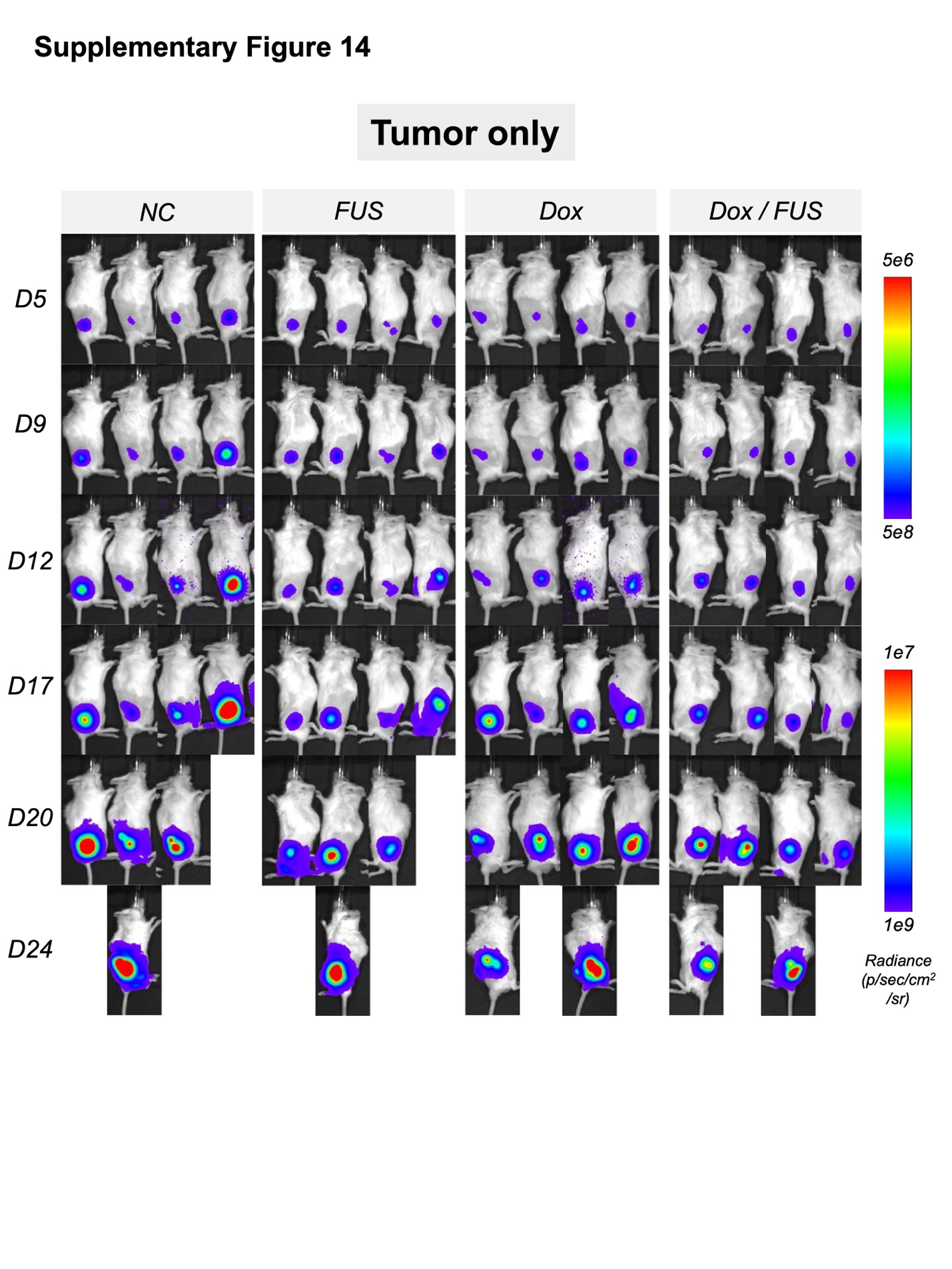

**Supplementary Figure 14: Effect of various treatments on tumor growths.** Bioluminescence (BLI) images showing tumor growth in a bilateral tumor model on mice (total 8 mice) without synNotch T cell administration. Tumors were activated on Day 10 (Fig. 6A) and tumor growth was monitored based on constitutive Fluc expression every three to four days starting from Day 5.

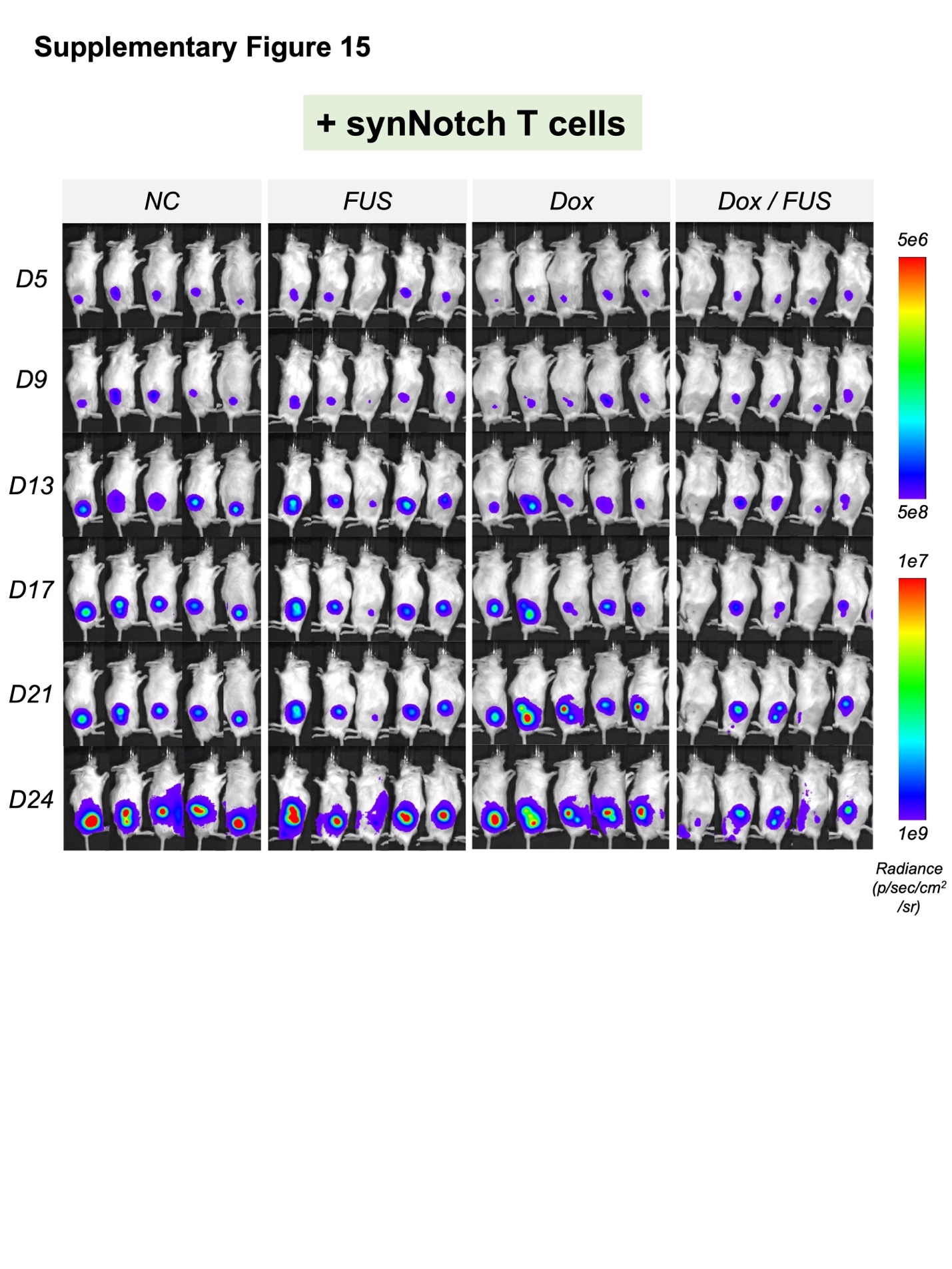

**Supplementary Figure 15: Effect of various treatments on tumor growths with synNotch CAR T cells.** Bioluminescence (BLI) images showing tumor growth in a bilateral tumor model on mice (total 10 mice) with synNotch CAR T cell administration. Tumors were activated on Day 10, followed by subcutaneous administration of synNotch CAR T cells. Tumor growth was monitored based on constitutive Fluc expression every three to four days starting from Day 5.

| Plasmids | Descriptions | Sources |
| --- | --- | --- |
| CMV-NFAT(4-460)-EGFP | NFAT translocation tracking | Addgene (#11107) |
| CMV-NFAT_Variants-EGFP | Kinetic study for NFAT variants | This study |
| PGK-rtTA-VIVIT_NFAT(4-399)-EGFP | CaDox regulator (EGFP) | This study |
| TRE-FLuc-PGK-RLuc-mCherry | CaDox reporter (FLuc/RLuc) | This study |
| TRE-mCherry | CaDox reporter (mCherry) | This study |
| PGK-rTetR-VIVIT_NFAT(4-399)-VP64-mCherry | CaDox regulator (mCherry) | This study |
| TRE-NLuc-PGK-EGFP | CaDox reporter (NLuc) | This study |
| CMV-R-Geco1 | Genetically encoded calcium sensor | Addgene (#32444) |
| TRE-tCD19-EGFP-BSD | CaDox reporter (tCD19) | This study |
| PGK-𝛼CD19_synNotch-GAL4-VP64 | Anti-CD19 synNotch construct | Addgene (#79125) |
| UAS-BFP-PGK-mCherry | Inducible BFP reporter | Addgene (#79130) |
| UAS-𝛼PSMA_CAR-PGK-mCherry | Inducible PSMA-CAR reporter | This study |

**Supplementary Table 1:** A list of constructs used in this study.

| Components | Final concentration |
| --- | --- |
| 50x B27 | 50x Diluted to make 1x |
| 500 mM N-Acetylcysteine | 1.25 mM |
| 0.5 mg/mL EGF | 5 ng/mL |
| 100 ug/mL Noggin | 100 ng/mL |
| R-Spondin 1 | 10% conditioned medium 500 nM |
| 5 mM A83-01 | 10 ng/mL 5 ng/mL 1 μM |
| 0.1 mg/mL FGF10 | 10 mM 10 μM 10% |
| 50 μg/mL FGF2 | 1 nM |
| 10 mM Prostaglandin E2 | 50x Diluted |
| 1M Nicotinamide | 1.25 mM |
| 30 mM SB202190 | 5 ng/mL |
| Fetal Bovine Serum | 100 ng/mL |
| 1000 mM HEPES | 10 mM |
| 200 mM GlutaMAX | 2 mM |
| 100x Pen-Strep | 1x |
| adDMEM/F12 |  |
| 100 mM Y-27632 Dihydrochloride | 10 µM |

**Supplementary Table 2:** The complete organoid media composition, including all components and their respective concentrations, was adapted from a previously published paper^45^ and is presented here.
